# RPFace: From biased structural benchmarks to generalizable RNA–protein interface prediction

**DOI:** 10.64898/2026.11.04.764332

**Authors:** Helyasadat Hashemi Aghdam, Sobhan Shukueian Tabrizi, A. Ercument Cicek

## Abstract

Modeling protein–RNA complexes with co-folding and docking methods is routine, yet these methods are evaluated on benchmarks whose composition mirrors the deposited structures on which they were trained or tuned, leaving open whether their reported performance reflects true generalization. Here, we focus the evaluation on the region that carries the biological signal: the binding interface. For the first time, we construct and release a fair benchmark that stratifies protein–RNA data into distinct structural and experimental groups, providing a more stringent test of generalization than aggregate averages. On this benchmark, we find that accuracy of detecting the binding site using co-folding methods is concentrated in ribosomal complexes and varies systematically with factors such as experimental modality. For docking methods, it is reliable only when the input conformations are already close to the bound state, and it is inconsistent across RNA classes. Both families of methods fail to transfer to *in vivo* CLIP experiments. To address this gap, we introduce RPFace, to the best of our knowledge the first model that jointly predicts the binding preference of every protein residue and every RNA nucleotide directly from sequence, requiring no experimental structure, template, or multiple sequence alignment. RPFace combines an ensemble of predicted secondary structures with a multi-modal embedding-enrichment scheme that fuses representations from RNA foundation models and structure prediction models. Unlike existing approaches, it is a very small and fast model that delivers stable performance across structural and experimental conditions, is robust to conformational change, and generalizes to *in vivo* datasets without extensive pretraining. As a result, RPFace matches the state-of-the-art methods on protein interfaces with no evolutionary information, and outperforms them on RNA interfaces, with the largest gains in zero-shot settings despite using only sequence information. The model, code, and data are available at Github.

## 1 Introduction

Protein–RNA interactions control translation, RNA processing, localization, and many other cellular processes. Identifying their binding sites is important for understanding how proteins recognize RNA and can guide further experimental studies. Experimental characterization of RNA–protein complexes can be demanding, which motivates computational approaches for predicting these interactions [1, 2]. However, predicting binding sites from molecular structures remains difficult because RNA is flexible and often undergoes conformational changes upon binding [3]. Moreover, available RNA–protein structural datasets are limited and strongly biased toward a few well-studied families, especially ribosomal complexes [4]. As a result, good average performance on structural benchmarks may not reflect generalization to less represented groups of proteins and RNAs.

Computational methods infer the interface by determining the three-dimensional structure of the complex. State-of-the-art co-folding models, such as AlphaFold3, Boltz-2, and RoseTTAFoldNA, first predict the tertiary structure of the complex and then extract the interacting residues from it [5–7]. These models have been shown to depend strongly on the evolutionary information provided by multiple sequence alignments (MSAs) [8]. Constructing such alignments requires extensive database searches, and informative homologs are often sparse or unavailable for poorly characterized or synthetic molecules [9]. Docking methods take a different approach: starting from experimentally determined or predicted structures of the individual molecules, they search for the binding pose that best positions and orients one partner relative to the other [10, 11, 13–16]. However, RNA–protein docking remains particularly challenging because the input structures change upon binding, and performance degrades sharply when the unbound conformations differ substantially from those in the bound complex [2, 17]. Another concern is that the reported accuracies of these tools are typically averaged over benchmarks whose composition closely mirrors the deposited structures on which the models were trained or tuned, so that a method may appear to perform well simply because the test set is dominated by the same families, folds, and experimental sources it has already seen [18, 19]. As a result, it remains unclear whether the published performance reflects a genuine generalization or a favorable overlap between training and evaluation data.

In this work we shift the focus of evaluation away from global structural accuracy and toward the region that actually carries the biological signal: the binding interface. The first contribution of this study is to release what is, to our knowledge, the first benchmark specifically designed to probe generalization in protein–RNA interface prediction. Rather than reporting a single aggregate score, it stratifies protein–RNA data into distinct structural and experimental groups, including homology, structure, RNA-family, complex, and time-based breaks, so that performance can be examined within each group and compared between them without leakage. Using this benchmark, we find that accuracy in detecting the binding site is concentrated in ribosomal complexes for co-folding methods, which are heavily represented in the training data. We also find that the performance varies systematically with factors such as the experimental technique used to solve the reference structure. For docking methods, binding-site prediction is reliable only when the supplied input conformations are already close to the bound state, and performance is inconsistent across RNA classes. Finally, both families of methods fail to transfer to *in vivo* CLIP experiments: co-folding models are trained on relatively stable, well-resolved PDB structures that have little resemblance to the cellular context captured by CLIP, while docking methods require three-dimensional input that such experiments cannot provide.

Our second contribution is RPFace, a lightweight, sequence-based model that predicts partner-specific binding preferences for protein–RNA pairs (Fig. 1). Given only the sequences of a protein and an RNA, RPFace assigns a binding propensity to every residue and every nucleotide, conditioning each side of the prediction on the identity of its partner. It requires no experimental or predicted structure, no template, and no MSA, which removes the dependence on structural databases and homology searches that limits co-folding and docking approaches on poorly characterized or synthetic molecules. The model trains only 2.1 million parameters, about 250 times fewer than Boltz-2, because it builds on frozen pretrained representations, and it predicts a pair in a median of 50 s from sequence alone. Ablation studies show that the frozen language-model views carry most of its accuracy (Section 3.4). On the 233-pair structural benchmark it reaches 0.88 AUROC on protein residues and 0.80 on RNA nucleotides, matching AlphaFold3 with alignments on proteins (0.87) and exceeding it on RNA (0.77). The gap widens exactly where structural data are thin. Without MSAs, AlphaFold3 falls to 0.64 and 0.52 while RPFace is unchanged by construction. Outside ribosomal complexes, AlphaFold3 drops to 0.60 on RNA against 0.76 for RPFace. On a January 2026 complex with a SELEX-derived RNA aptamer, which has no homologs at all, RPFace reaches 0.855 RNA AUROC against 0.694 for the best co-folding configuration. On a docking benchmark excluded from our training data, RPFace obtains the highest protein AUROC (0.782) from sequence alone, against docking methods supplied with the unbound structures, and its accuracy varies by 0.003 across difficulty classes where the best docking method loses 0.16. We further test whether interface prediction learned from structural data transfers to the cellular context by fine-tuning and evaluating RPFace on *in vivo* CLIP data from CLIPdb [20, 21]. The model generalizes to cell lines and chromosomes held out during training, while the drop relative to the structural benchmark exposes a persistent gap between the interface in a solved complex and RNA binding as it occurs in the cell. The model, code, and benchmark data are available at Github.

**Figure 1:**
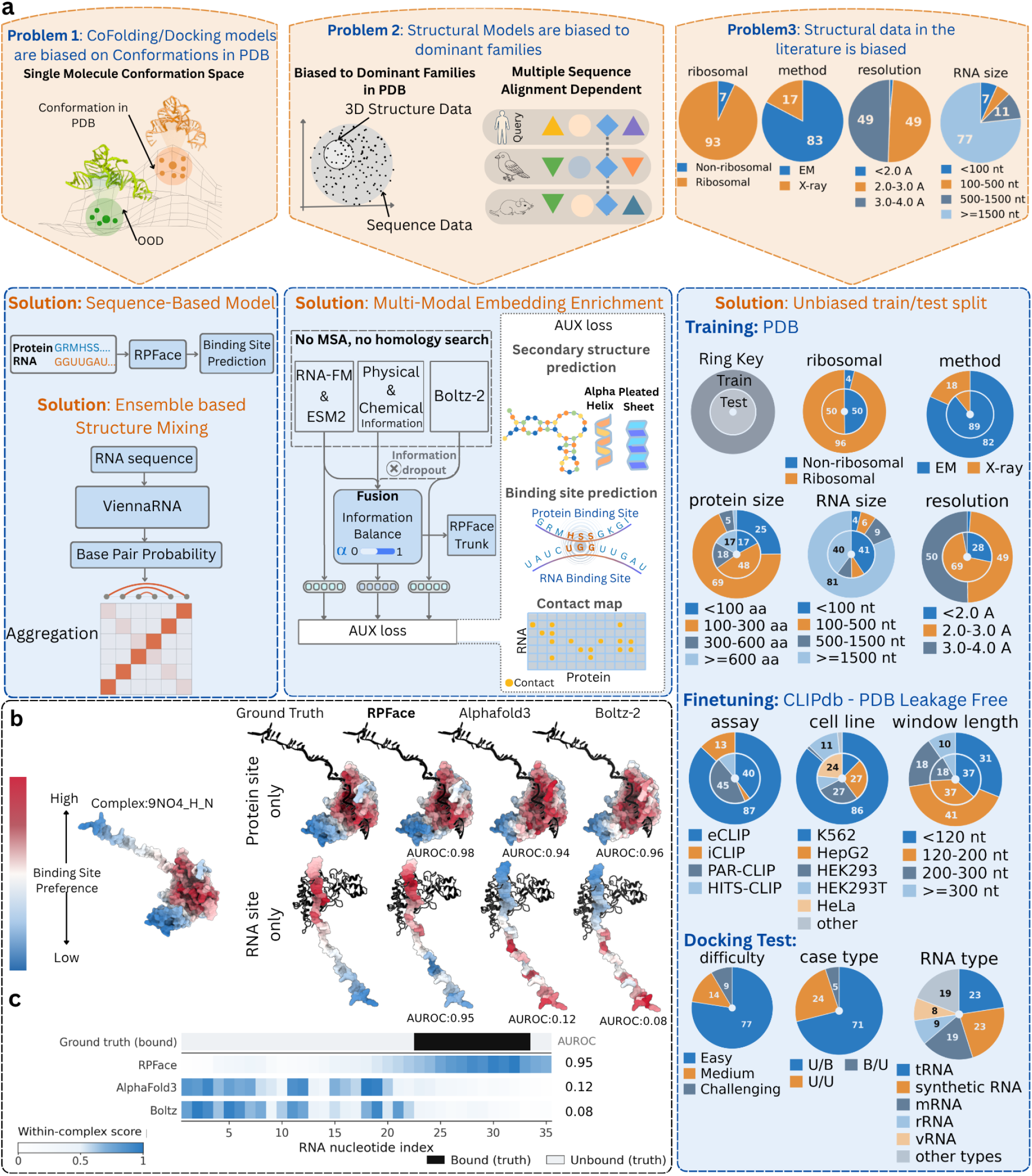
Biases in protein–RNA structural data and how RPFace addresses them. (**a**) Three problems (top) and the corresponding solutions (bottom). (1) Co-folding and docking methods are biased towards conformations deposited in the PDB. RPFace uses sequences alone and aggregates ViennaRNA base-pair probabilities over the RNA ensemble. (2) Structural models are biased towards PDB families and depend on MSAs. RPFace fuses language-model embeddings (ESM2, RNA-FM), physicochemical descriptors and an MSA-free Boltz-2 embedding through a learned gate (*α*), with auxiliary losses used only in training. (3) PDB structures are dominated by a few groups. We use leakage-free splits and evaluate each group separately (PDB panel: outer rings are the training set, inner rings the 233-pair benchmark, with values in percent). Fine-tuning data come from CLIPdb, and docking tests use the benchmark of Chong et al. [22]. (**b**) Predicted binding preference for PDB 9NO4, from high (red) to low (blue), with protein at top, RNA at bottom and partner in black. (**c**) Per-nucleotide RNA scores for the complex in **b**, with the ground truth on top (black, bound) and each method’s AUROC on the right.

## 2 Methods

Interface predictors are usually scored by a single average over a test set drawn from the same distribution as the training data. Because protein–RNA structures are dominated by a few families, such an average mostly reports performance on those families. Our benchmark is built so that performance can be read separately within each structural and experimental group, and so that no group leaks between training and evaluation. We built complementary structural and *in vivo* datasets (Table 1) from protein–RNA structures in the PDB [23] and human CLIP peaks in POSTAR3 [21]. Full procedures are given in Supplementary Sections S1 and S2.

**Table 1:**
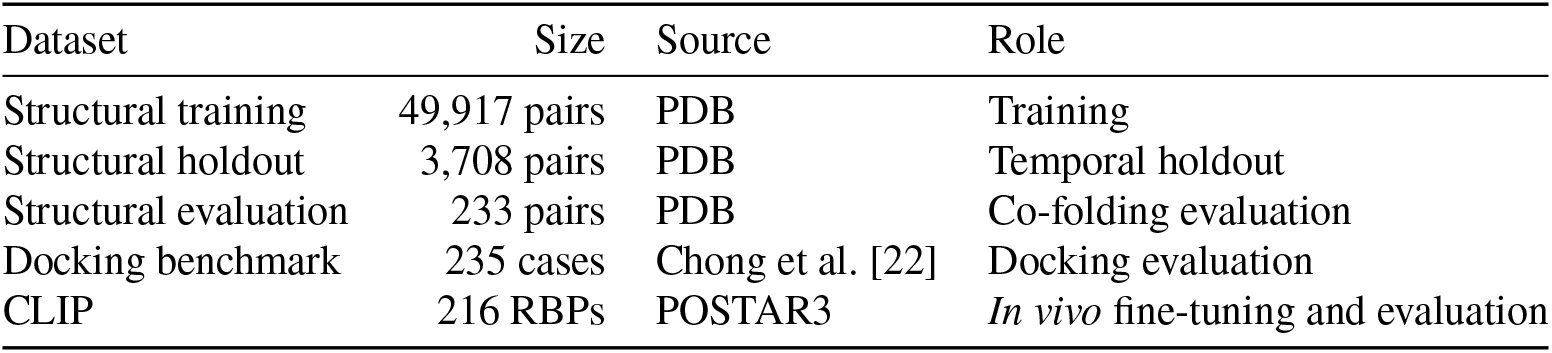
Datasets. The 233-pair structural evaluation subset is drawn from the temporal holdout and contains 117 ribosomal and 116 non-ribosomal pairs, and 208 cryo-EM and 25 X-ray structures. RBP, RNA-binding protein.

| Dataset | Size | Source | Role |
|---|---|---|---|
| Structural training | 49,917 pairs | PDB | Training |
| Structural holdout | 3,708 pairs | PDB | Temporal holdout |
| Structural evaluation | 233 pairs | PDB | Co-folding evaluation |
| Docking benchmark | 235 cases | Chong et al. [22] | Docking evaluation |
| CLIP | 216 RBPs | POSTAR3 | *In vivo* fine-tuning and evaluation |

### 2.1 Structural benchmark

#### Structural data

We selected PDB entries containing protein and RNA with resolution ≤ 3.5 Å (biological assembly 1). For each protein–RNA chain pair we computed all heavy-atom distances within the complete assembly, and kept pairs with at least 25 inter-chain contacts within 4.5 Å. Deduplication on sequence and interface signatures gave 67,716 candidate pairs.

#### Leakage control against the docking benchmark

Docking evaluation needs both bound and unbound structures, which are rare, so we used the 235-case benchmark of Chong et al. [22] rather than assembling our own. Any training pair resembling one of its cases would invalidate that comparison, so we removed related pairs in four steps before clustering (Table 2). First we removed pairs from its PDB entries. Second we removed proteins with ≥30% identity over ≥50% coverage to its proteins (MMseqs2 [24]). Third we removed RNAs assigned to one of its Rfam families (Infernal [25, 26]). Finally we removed proteins with a Foldseek match of max (qTM, tTM) ≥0.5 [27]. Structural homology removed by far the most data, 9,376 pairs after sequence homology had already been applied, which shows that sequence-identity filtering alone would have left substantial overlap. The four steps together left 53,625 pairs.

**Table 2:** Leakage filtering of the structural data against the docking benchmark. Counts are protein–RNA chain pairs. Each step is applied to the output of the previous one.

| Step | Removed | Remaining |
|---|---|---|
| Initial deduplicated pool | – | 67,716 |
| PDB entries of the docking benchmark | 216 | 67,500 |
| Protein sequence homology (MMseqs2) | 2,783 | 64,717 |
| Rfam families of benchmark RNAs (Infernal) | 1,716 | 63,001 |
| Protein structural homology (Foldseek) | 9,376 | 53,625 |

#### Clusters and the temporal split

We clustered the filtered set once, obtaining 1,098 protein and 635 RNA clusters, and defined a complex cluster as the combination of a protein cluster and an RNA cluster, giving 9,029 complex clusters. The same protein bound to a different RNA, or the same RNA bound to a different protein, therefore falls in a different cluster. This product definition is deliberate: a predictor that is conditioned on both partners should be tested on partner combinations it has not seen, not merely on unseen molecules. We used June 1, 2024 (the Boltz-2 training cutoff) as the temporal cutoff, so that the evaluation structures postdate the training data of the co-folding models we compare against as well as our own. Only clusters whose structures all postdate the cutoff were eligible for the holdout. A cluster containing any earlier structure was assigned entirely to training. No complex cluster is shared between the two partitions. The validation set is 10–15% of the eligible clusters, sampled with stratification by experimental method, ribosomal status, resolution (< 2.5, 2.5–3.0, ≥ 3.0 Å) and RNA length (< 20, 20–50, 50–100, > 100 nt), so that it does not collapse onto a single regime. For the co-folding comparison, we drew a fixed 233-pair evaluation subset from the holdout, balanced between ribosomal and non-ribosomal complexes (117 and 116) so that the stratified comparison in Section 3.1 has comparable support on both sides.

#### Composition and sampling

The remaining imbalance is severe: ribosomal complexes supply about 90% of the interface-centred training crops, and the largest 1% of clusters alone contribute 20% of them. Training therefore draws crops with cluster weights proportional to (1/cluster size) ^0.5^ within the ribosomal and non-ribosomal groups separately, and reweights the two groups from 90:10 to 70:30, matching the validation set. This raises the effective number of clusters from 2,575 to 5,401 and lowers the share of the largest 1% of clusters from 20% to 7% (Supplementary Fig. S1b).

### 2.2 CLIP benchmark

#### CLIP data

POSTAR3 provides 38.4 million human peaks on GRCh38, which we grouped by CLIP technology and cell line or tissue. We kept groups with at least two proteins and 5,000 peaks, and proteins with at least 200 peaks in a group. Of the 52 groups, 16 qualified, spanning eCLIP, PAR-CLIP, iCLIP and HITS-CLIP. Requiring two proteins per group supplies cross-protein negative evidence: a nucleotide bound by another protein in the same experimental context is a more informative negative than an unassayed one. Peak scores from different callers measure different quantities and are not comparable, so we did not use them.

Within each group, overlapping peaks were merged per protein, and the merged peaks of all proteins were combined into contiguous coverage blocks. Windows were sampled only from inside these blocks, at most three per block, so that every included nucleotide is covered by at least one experiment in the same context. The RNA sequence of a window is the corresponding genomic sequence from the GRCh38 primary assembly. Gene annotations come from GENCODE v29 and full-length RBP sequences from UniProt [12]. Each window is assigned an anchor protein, which must retain at least 50% of its peak after windowing and cover 10–45% of the window. Among eligible proteins we chose the least-used one, so that frequent proteins and densely bound loci do not dominate. A nucleotide is positive if it lies in a merged peak of that protein, and a window–protein pair is discarded if a non-anchor peak is truncated below 15 nt by a window boundary. This yields windows of 50–1,450 nt across 216 RBPs.

#### CLIP leakage control and evaluation axes

Because CLIP fine-tuning starts from the structurally trained model, the two datasets were protected jointly, in both directions, at the same thresholds as above. Of the 216 CLIP RBPs, 190 were unrestricted, 14 were homologous to proteins in the structural evaluation set and were excluded from CLIP training, 5 were homologous to structural training proteins and were excluded from the CLIP holdout, and 7 matched both partitions and were dropped entirely. Restrictions were applied at the level of the 174 homology clusters these proteins form, not to individual proteins. On the RNA side, windows overlapping structured non-coding RNA classes or Rfam families common in the PDB were removed. Three axes were then held out independently: assay (PAR-CLIP, HITS-CLIP), cell line (HepG2, HeLa) and chromosome (chr1, chr8). Only the chromosome axis guarantees unseen RNA sequence, since a locus held out by assay or cell line may still appear in another retained experiment. We therefore report a *strict* version of the assay and cell-line slices restricted to the held-out chromosomes, alongside a *relaxed* version without that restriction.

### 2.3 Graded interface targets

Given a protein–RNA pair (***a, b***) with *P* residues and *R* nucleotides, RPFace predicts a continuous binding preference for every residue and nucleotide. A position belongs to the interface if its minimum heavy-atom distance to the partner is at most 4.5 Å. Binary interface labels are strongly imbalanced, so we supervise with graded targets that decrease with the distance from the interface. Let 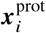 be the mean heavy-atom coordinate of residue *i* and ℐ^prot^ the set of protein interface positions. The distance to the nearest interface position on the same molecule, and the target, are

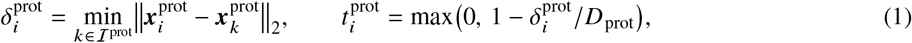

and analogously 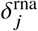 and 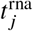 for RNA, with *D*_prot_ = 10 Å and *D*_rna_ = 30 Å. Interface positions receive 1, and nearby positions receive smaller values that reach zero at the cutoff. As a result, 36.6% of residues and 39.0% of nucleotides receive non-zero targets (Supplementary Fig. S1a). Complexes without contacts (spatial negatives) receive ***t*** ≡ 0. Because the target is an affine function of *δ* truncated to [0, 1], a prediction 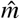 maps back to a distance 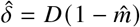 within the uncensored range. The model thus performs censored regression of the distance to the interface through a logistic link. The cutoffs are absolute rather than normalized per structure, which makes the targets independent of the geometry of the interface-centred training crops (Supplementary Section S3).

### 2.4 Input representations and gated fusion

Each position is described by three views. For proteins these are the Boltz-2 trunk representation ***s***^prot^ ∈ ℝ^*P*×384^, computed in single-sequence mode with an empty MSA [6], the frozen ESM2 embedding ***e*** ∈ ℝ^*P*×1280^ [28], and a prior ***q***^prot^. For RNA, they are the Boltz-2 representation ***s***^rna^, the frozen RNA-FM embedding ***g*** ∈ ℝ^*R*×640^ [29] and a prior ***q***^rna^. The Boltz-2 view contributes folding-based structural information, while the language models carry patterns learned from large sequence collections. The prior of token *a*_*i*_ is *q*_*i*_ = *ψ* ([Emb (*a*_*i*_); ***ϕ*** (*a*_*i*_); PE (*i*)]), a two-layer perceptron over a learned 32-dimensional token embedding, physicochemical descriptors ***ϕ*** (hydropathy, charge, aromaticity and volume for residues, and a purine indicator with hydrogen-bond donor and acceptor counts for nucleotides), and a 16-dimensional sinusoidal positional encoding. The descriptors are standardized with fixed population constants rather than batch statistics, so the encoding of a token does not depend on the complex in which it occurs. We project the structural, language-model and prior views to *d* = 96 dimensions with layer-normalized affine maps, giving *h*_1,*i*_, *h*_2,*i*_ and *h*_0,*i*_, and combine them with a per-token convex gate

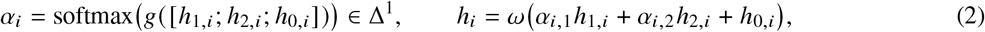

where *g* is a two-layer gating network and *ω* an output projection. The prior enters as an ungated residual that the gate cannot remove. Because *α*_*i*_ is computed per token, the balance between structural and language-model information is set locally rather than once per complex. During training, the projected Boltz-2 view is dropped with probability 0.1. For CLIP fine-tuning, we remove the Boltz-2 view entirely (Section 2.8).

### 2.5 Ensemble-averaged secondary-structure propagation

RNA in a solved complex is one of many conformations the molecule adopts. Let ***M*** ∈ [0, 1] ^*R*×*R*^ be the McCaskill base-pairing probability matrix of ***b***, computed with ViennaRNA under the Turner energy model [30]. We symmetrize and row-normalize it, 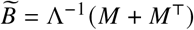 with Λ = diag(∑_*j*_ (***M*** + ***M***^⊤^)_*ij*_), and apply one gated propagation step

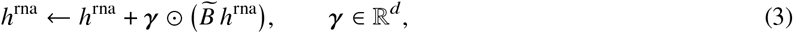

with the per-channel gate ***γ*** initialized at 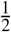. The term 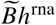 is the expected representation of a nucleotide’s pairing partner under the equilibrium Boltzmann ensemble, not the single partner of a minimum-free-energy or crystallographic structure. It therefore does not depend on any particular conformer. This is the only message-passing step in the final model: ablations supported removing three alternatives, namely local-sequence propagation on the protein and on the RNA, and propagation over RNA-FM embedding similarity.

### 2.6 Partner coupling and multi-scale mixing

The protein and RNA tracks are coupled by bidirectional cross-attention with four heads, so each residue attends to the RNA and each nucleotide to the protein. A compact pair representation is updated from both tracks. A second per-token convex gate *β*_*i*_ mixes the representations before (*h*_loc_) and after (*h*_crs_) cross-attention, letting the model down-weight partner information when it is uninformative. Neither the Boltz-2 pair representation nor predicted coordinates are used. Both tracks are then processed by a masked dilated convolutional stack (dilations 1 to 32, kernel width 5, gated linear units, residual connections) with a receptive field of 253 positions. This captures extended binding regions without a fixed input window and supports transfer from interface-centred training crops to full-length chains at inference. Each position is finally mapped to a single binding score.

### 2.7 Training objective

The main loss is the cross-entropy between predicted logits *u*_*i*_, *v* _*j*_ and the graded targets, averaged over the valid positions ℳ of each molecule pooled across the batch:

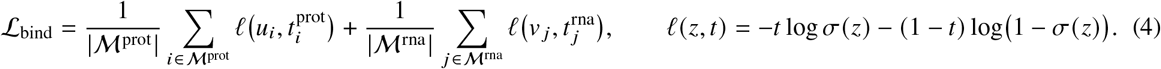

Its unique minimizer is *σ* (*z*) = *t*, so it is proper for recovering the graded profile. Its attainable minimum is the target entropy, which is non-zero only on the graded rim around the interface. Because *D*_rna_ = 3*D*_prot_, the RNA term has a higher floor, so protein and RNA loss values are not directly comparable. Cross-entropy treats positions independently and is dominated by long chains. We therefore add a per-sequence Lovász hinge loss ℒ_lov_ on the binary interface labels, a convex surrogate of the Jaccard loss that directly rewards set-level overlap. Three auxiliary terms, used only during training, supervise a latent protein–RNA contact map from the pair representation (ℒ _pair_, against a graded inter-chain contact map built as in Eq. (1)), RNA base pairing (ℒ _ss_) and three-state protein secondary structure (ℒ _sse_). To prevent shortcut learning, we attach binding heads to each projected view (ℒ_view_) and to the representation before partner interaction (ℒ _pre_). Together with view dropout, these heads are intended to keep the gate from collapsing onto whichever view is easiest to optimize early in training; removing the auxiliary losses shifts the protein gate toward the Boltz-2 view without changing accuracy (Supplementary Section S9). Finally, the regularizer 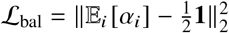 balances the batch-averaged gate while leaving individual tokens free to commit to one view. The total loss is ℒ = ℒ_bind_+ ∑_*k*_*λ*_*k*_ ℒ_*k*_ over these terms, with *λ*_lov_ = *λ*_pair_ = *λ*_pre_ = 0.1, *λ*_sse_ = *λ*_view_ = 0.2, *λ*_ss_ = 1 and *λ*_bal_ = 0.01. The Lovász weight was selected on the validation set, and all other weights were fixed before training (Supplementary Section S4). Training uses interface-centred crops, including spatial negatives, drawn by the cluster-balanced sampler of Section 2.1.

#### Implementation details

RPFace trains 2.1 million parameters on top of the frozen ESM2, RNA-FM and Boltz-2 models; Boltz-2 representations are precomputed. We trained for 200 epochs with AdamW (learning rate 10^−3^, weight decay 10^−2^), linear warm-up over the first 10% of steps followed by cosine decay, global gradient-norm clipping at 1.0 and random seed 0.

### 2.8 CLIP fine-tuning

RPFace-CLIP is initialized from the structural model and adds no parameters: CLIP peaks supervise the same RNA binding head, and the Boltz-2 view is dropped from the fusion of Eq. (2), so every CLIP input is derived from sequence. Structural supervision is kept throughout. Each optimizer step combines one structural and one CLIP batch, minimizing

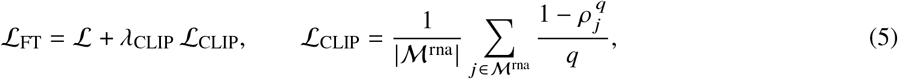

Where ℒ is the structural objective above and *ρ* _*j*_ is the predicted probability of the observed peak label of nucleotide *j*. Because a single head must serve both label types, the structural term is what holds it to the interface task. ℒ_CLIP_ is the generalized cross-entropy [31] with *q* = 0.7. It down-weights positions whose label the model confidently contradicts, which suits the one-sided noise of peak calls. We set *λ*_CLIP_ = 0.114, at which the gradient norms of the two terms in the shared head are equal; without weighting, the CLIP term supplies about 91% of the update. CLIP labels exist only for RNA, so the protein head is trained by the structural term alone. Fine-tuning uses AdamW with a learning rate of 10^−4^ and no weight decay, the same schedule and clipping as above, and batches of 16 windows, reduced for long inputs to bound GPU memory.

### 2.9 Evaluation

Because RPFace outputs continuous preferences, we evaluate rankings rather than thresholded contacts. Experimental contacts define the positives. Structure-based methods are scored from their predicted coordinates as *s* = 1/*d*, where *d* is the minimum distance from a residue or nucleotide to the partner molecule. All methods therefore produce continuous rankings without method-specific thresholds. We report AUROC, AUPRC, Precision@*K* and Success@5 (Supplementary Section S5). For Precision@*K, K* is the number of interface positions of the molecule, so it equals Recall@*K* and a perfect ranking scores 1. Success@5 indicates whether any of the five top-ranked positions is an interface position. AUROC does not depend on prevalence, whereas AUPRC, Precision@*K* and Success@5 are read against the interface fraction of each stratum. Intervals are 95% bootstrap intervals that resample complex clusters on the PDB benchmark and complexes on the docking benchmark.

For CLIP data, standard ranking metrics are biased by the width of the called peaks, so we use a dedicated enrichment metric, M3, defined in Section 2.10. Chance is measured rather than assumed. For M3, a circular-shift null preserves each score profile’s distribution and autocorrelation and destroys only its alignment to the peaks. A positional baseline scores only proximity to the window centre, which a model must exceed to show sequence-specific signal. Intervals are bootstraps over RBPs rather than windows, since windows of one protein are correlated, and methods are compared by paired per-RBP differences.

### 2.10 M3: a peak-width-robust metric for CLIP data

CLIP peaks mark crosslinked fragments that are usually much wider than the protein footprint, and ranking metrics computed against them penalize a predictor that finds the true site (Section 3.3). We therefore introduce normalized mass enrichment (M3), which measures where a prediction places its score mass rather than how it ranks individual nucleotides. For a window of *L* nucleotides with peak labels *y*_*i*_ and coverage *p* = ∑_*i*_ *y*_*i*_/*L*, we first remove the baseline of the score profile, 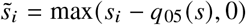, where *q*_05_ is its fifth percentile, so that a uniform score floor does not dilute the mass. We then compute

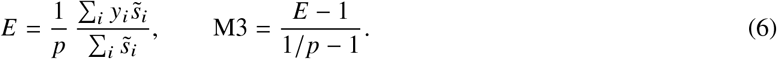

*E* is the share of score mass inside the peak relative to the share expected by chance. M3 has four properties that standard metrics lack on CLIP data. (i) *Calibrated chance level*. For any coverage *p*, mass unrelated to the peak gives 0 in expectation and full concentration inside it gives 1. Values lie in [−*p*/(1 – *p*), 1], and negative values mark mass depleted from the peak. (ii) *Robustness to peak dilation*. Concentration anywhere inside the peak counts, so a predictor that marks only a footprint narrower than the peak still reaches 1. (iii) *Comparability across windows*. Normalizing by *p* removes the dependence on peak coverage that makes AUPRC and P@*K* incomparable between windows. (iv) *Independence of score units*. M3 is invariant to positive affine transformations of the scores, so it does not depend on the scale or offset of a method’s output. Unlike AUROC, it is not invariant to non-linear monotone transformations, so it reflects the shape of the score profile and not only its ranking. An M3 value converts back to an absolute share of mass inside the peak, *p* + M3 (1 − *p*). At the median coverage *p* = 0.29, an M3 of 0.1 means that 36% of the mass lies inside the peak, compared with 29% at chance. Constant profiles, for which 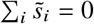, are excluded. Chance is measured with the circular-shift null of Section 2.9 rather than assumed. Proofs of the properties are given in Supplementary Section S5.

## 3 Results

### 3.1 RPFace generalizes beyond evolutionary information and the training distribution

#### Dependence on MSAs

We compared RPFace with AlphaFold3 [5] and Boltz-2 [6], each run with and without MSAs, on the 233-pair evaluation subset (Table 1). Removing MSAs substantially reduces co-folding performance (Fig. 2a). On proteins, AlphaFold3 drops from 0.87 to 0.64 AUROC and Boltz-2 from 0.87 to 0.69, whereas RPFace, which never uses MSAs, reaches 0.88. On RNA the drop is larger: AlphaFold3 falls from 0.77 to 0.52 and Boltz-2 from 0.67 to 0.56, while RPFace achieves 0.80.

**Figure 2:**
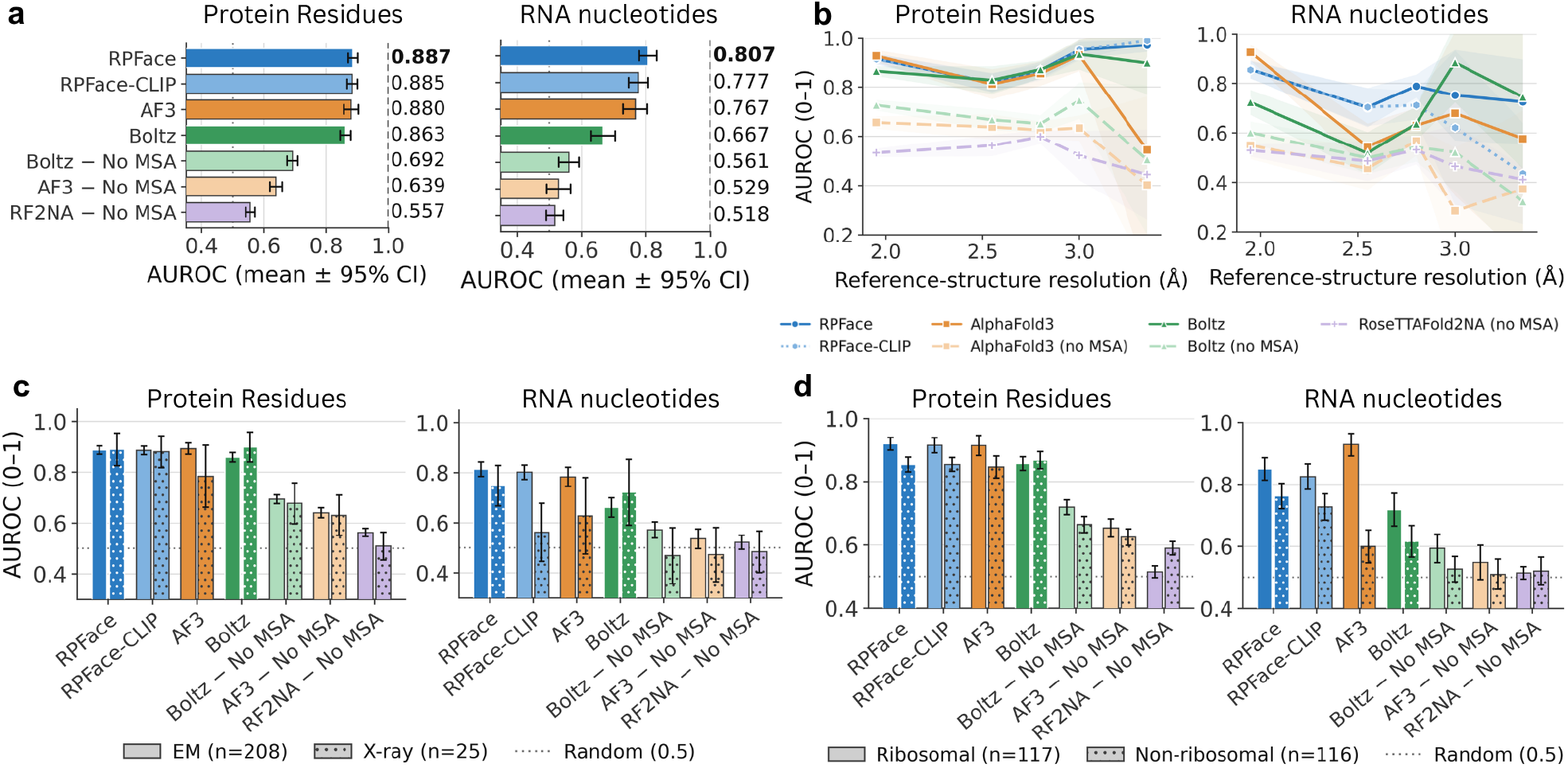
Comparison with co-folding models on the 233-pair structural benchmark. **a** Mean AUROC (± 95% CI) for protein residues (left) and RNA nucleotides (right); removing MSAs causes a large drop for Boltz-2 and AlphaFold3. **b** AUROC by reference-structure resolution; RPFace remains consistently high, while other methods are lower and more variable. **c** AUROC by structure-solution method: cryo-EM (*n* = 208) vs X-ray (*n* = 25). **d** AUROC by complex type: ribosomal (*n* = 117) vs non-ribosomal (*n* = 116).

#### Dominant data regime

Ribosomal complexes dominate the structural data (Supplementary Fig. S1b), and co-folding accuracy on RNA follows this bias. AlphaFold3 reaches 0.93 AUROC on ribosomal complexes but only 0.60 on non-ribosomal ones (Fig. 2d). RPFace drops by only 0.08 and is 0.16 higher than AlphaFold3 outside the ribosomal group. Boltz-2 drops by a similar amount from a much lower level. On proteins the effect is weak: all methods that use MSAs lose a few points and stay close to each other. The same pattern appears across experimental methods, where AlphaFold3 falls to 0.56 on X-ray RNA while RPFace keeps 0.75 (Fig. 2c). It also appears across resolution bins (Fig. 2b). Cryo-EM, low resolution and ribosomal identity are strongly correlated in the data, so we treat these as views of one distribution shift rather than independent effects.

#### An RNA partner without homologs

The MSA results suggest that co-folding models should struggle when a partner has no homologs. PDB 9I0V [32] tests this directly. It is a January 2026 structure of the *Streptomyces coelicolor* repressor DasR bound to a SELEX-derived RNA aptamer, released after both co-folding cutoffs. We identified it among post-cutoff entries whose RNA is annotated as a synthetic construct, and present it as a case study rather than a benchmark. DasR belongs to the widespread GntR/HutC family and has abundant homologs. The *in vitro*-selected aptamer has none, so it provides no intra-RNA or cross-chain coevolutionary signal.

On the RNA side (Fig. 3), RPFace reaches 0.855 AUROC and 0.875 AUPRC (contact prevalence 0.47). AlphaFold3 reaches 0.498 AUROC with alignments and 0.694 without them, and Boltz-2 reaches 0.431 in both settings. RPFace thus exceeds the best co-folding configuration by 0.161 AUROC and 0.253 AUPRC. On the protein side, alignments help the co-folding models strongly. With alignments, AlphaFold3 and Boltz-2 reach 0.973 and 0.967 AUROC (AUPRC 0.723 and 0.641) and outperform RPFace. Without alignments, both fall to the level of RPFace (0.811 and 0.797 vs. 0.786 AUROC; 0.184 and 0.177 vs. 0.179 AUPRC). The protein-side advantage of co-folding therefore derives largely from alignments. On the RNA side, where alignments carry no information, RPFace gives by far the strongest ranking.

**Figure 3:**
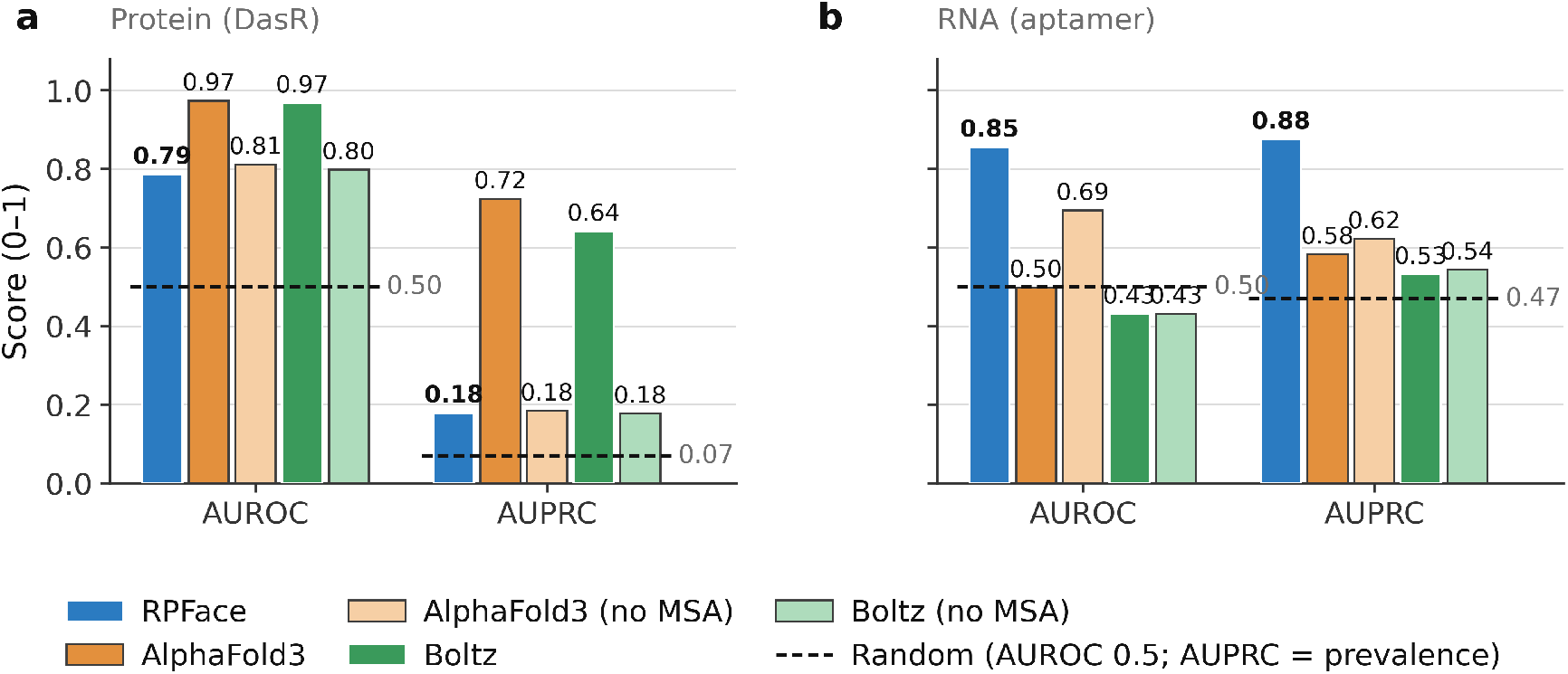
Interface prediction on PDB 9I0V (DasR bound to a SELEX-derived RNA aptamer). RPFace is compared with AlphaFold3 and Boltz-2, each run with and without MSAs. Left: protein side, where evolutionary information is available. Right: RNA side, where it is not. Dashed lines show chance (AUROC) and contact prevalence (AUPRC).

#### Comparison with a sequence-based method

We also compared RPFace with ProRB, a recent sequence-based predictor, on the FoldBench protein–RNA targets used in its evaluation [33, 34]. On the four targets independent of the RPFace training data, RPFace reached 0.708 protein and 0.715 RNA AUROC. The values reported for ProRB are

0.575 and 0.544, and for AlphaFold3 0.557 and 0.643. Those published values average over all nine targets, each of which has a close homolog in the ProRB training data. The overlap analysis and results on all nine targets are given in Supplementary Section S8.

Together, these results show that the interface accuracy of co-folding models depends on evolutionary information and on similarity to the dominant training regime, whereas RPFace remains stable when either is missing.

### 3.2 RPFace does not depend on the input conformation

On the docking benchmark of Chong et al. [22], whose complexes were excluded from our training data (Section 2.1), docking methods receive the unbound protein and RNA structures. RPFace sees only the two sequences, so the comparison favors docking. Even so, RPFace has the highest protein AUROC (0.782) and is level with HDOCKlite on RNA (0.644 vs. 0.645; Fig. 4a). The main difference is consistency. Across RNA classes, RPFace stays above chance everywhere and has the best class average. HDOCKlite is best on rRNA but falls below chance on snRNA and vRNA (Fig. 4d). The same pattern appears when targets are grouped by the conformational change between unbound and bound structures (Fig. 4b). Docking accuracy drops as the inputs move away from the bound conformation: HDOCKlite loses 0.16 protein AUROC between the easiest and hardest groups. RPFace, whose sequence input does not change with difficulty, varies by only 0.003.

**Figure 4:**
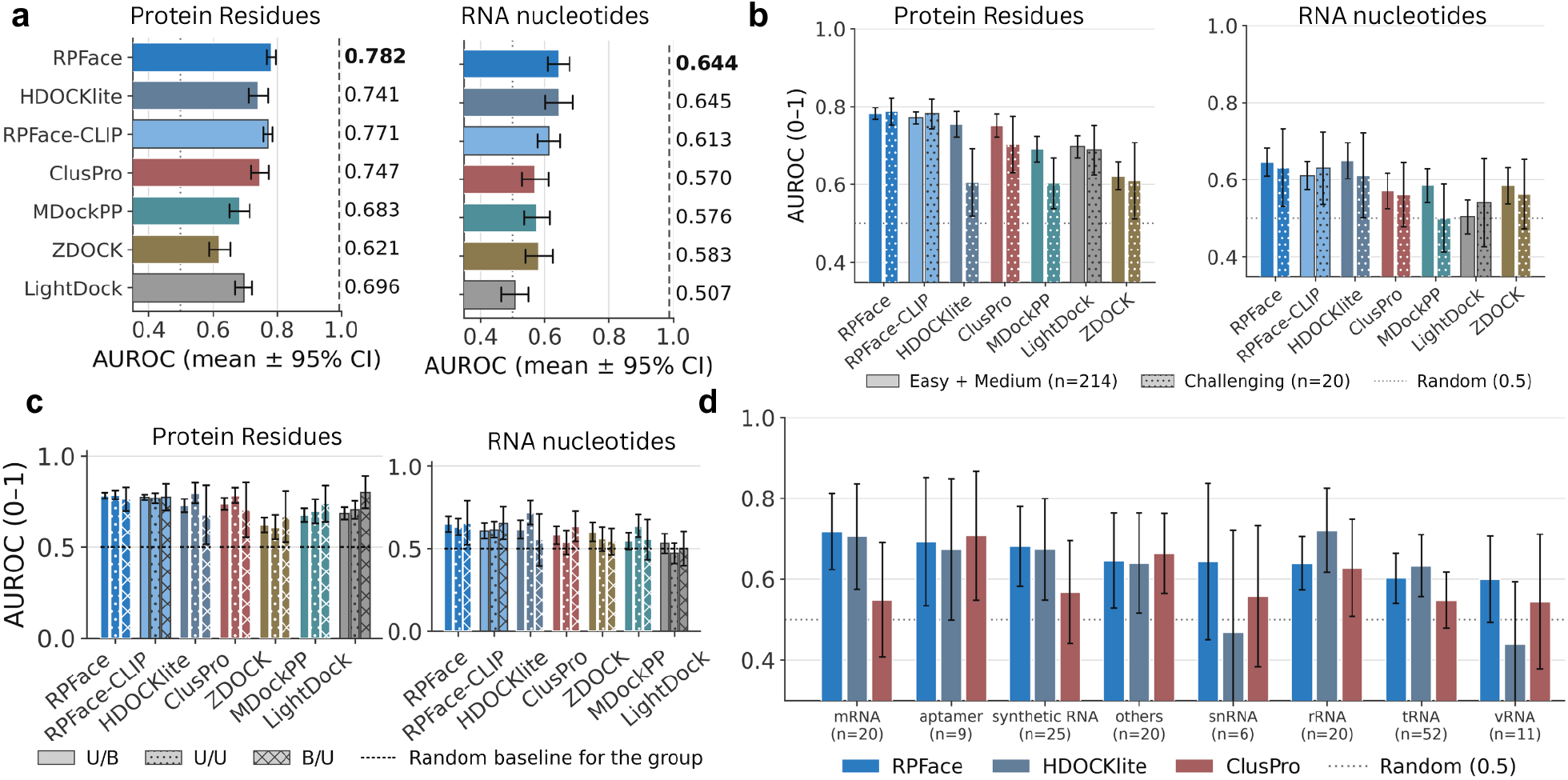
Comparison with docking methods. Docking methods are given the unbound structures, while RPFace predicts from sequence alone. **a** Overall AUROC on protein residues and RNA nucleotides (mean ±95% CI), with the number of scored cases beside each method. **b** AUROC with targets grouped by docking difficulty. **c** AUROC on protein residues (left) and RNA nucleotides (right), with targets grouped by the state of the input structures supplied to the docking methods: U/B, unbound protein with bound RNA; U/U, both unbound; B/U, bound protein with unbound RNA. RPFace and RPFace-CLIP receive only sequences, so their input is the same in every group; differences between their bars reflect the targets in each group. Bars show means with 95% CIs. **d** RNA nucleotide AUROC by RNA class. Dotted lines mark random expectation (0.5).

### 3.3 CLIP supervision enables transfer to *in vivo* binding

We next ask whether interaction scores learned from structures transfer to protein binding measured *in vivo*.

#### Limitations of standard metrics

A CLIP peak marks a crosslinked RNA fragment and is usually much wider than the protein’s physical footprint. Its positive labels therefore cover the footprint plus flanking nucleotides that the protein does not contact. Ranking metrics then penalize a predictor that localizes the true site correctly. Consider an oracle that scores a footprint of width *w* inside a peak of width *m* high and everything else uniformly low. Footprint positions outrank every negative and the remaining peak positions tie with them, so its AUROC is 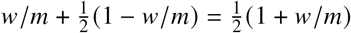, which falls toward chance as the peak widens (Fig. 5a). For an 8-nt footprint at typical peak widths, this ceiling is about 0.62. The oracle indeed obtains AUROC 0.62 and P@*K* 0.50 despite identifying the correct site (Fig. 5b). P@*K* falls because, once the footprint is exhausted, the remaining top-ranked positions are no better than chance. An oracle that simply reproduces the whole peak reaches 1.00 on both metrics. AUROC and P@*K* therefore reward matching the annotated peak width rather than finding the binding site. In addition, AUPRC and P@*K* depend on the fraction of a window covered by peaks. This fraction varies widely across windows, so their values are not comparable from one window to another.

**Figure 5:**
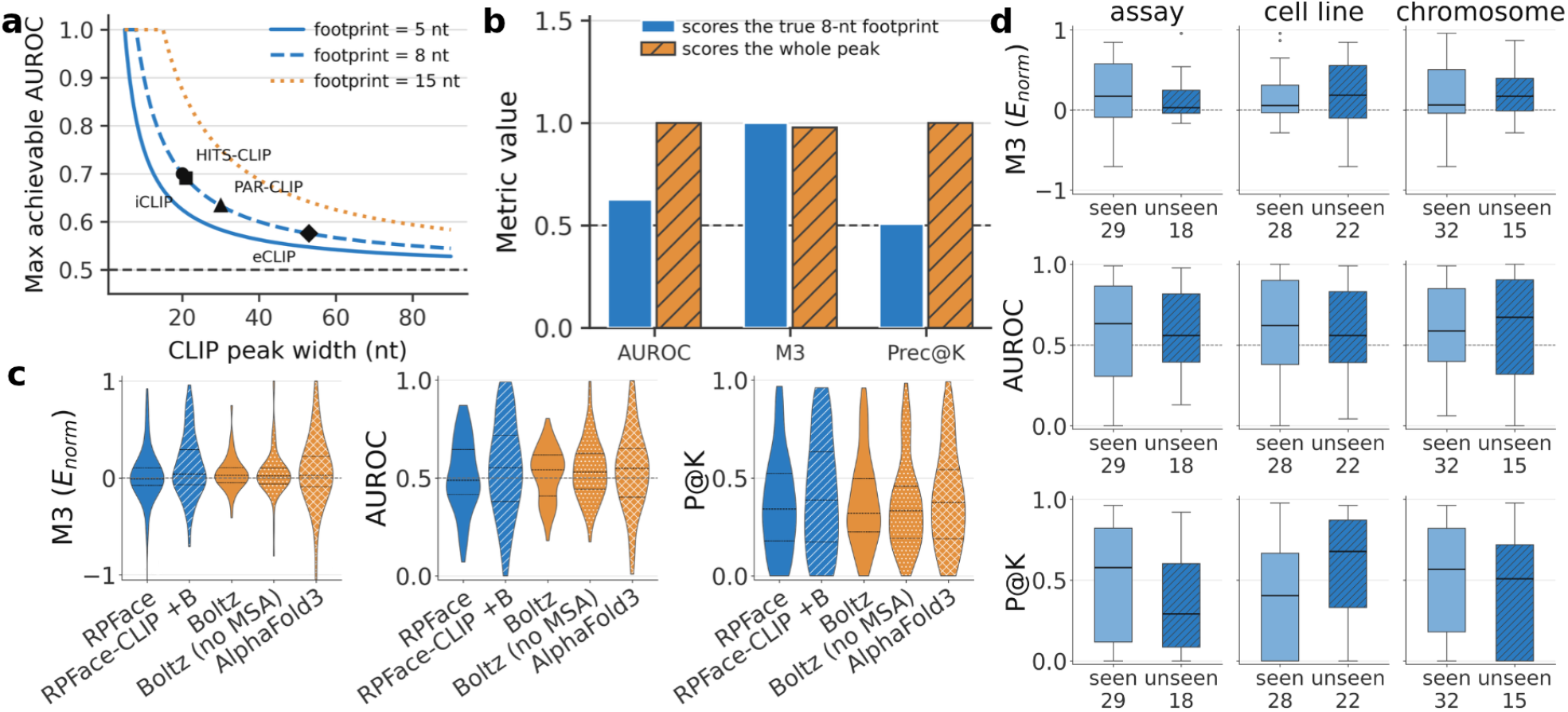
CLIP supervision enables *in vivo* binding prediction, whereas structural scores alone show little transfer. **a** Maximum achievable AUROC for a predictor that correctly identifies a footprint narrower than the annotated peak, for footprints of 5, 8 and 15 nt, as a function of peak width. **b** AUROC, P@*K* and M3 for two oracles: one scores only the true 8-nt footprint, and the other scores the whole peak. **c** RPFace, CLIP-fine-tuned RPFace (RPFace-CLIP), Boltz-2 with and without MSAs, and AlphaFold3 on the CLIP benchmark (M3, with AUROC and P@*K* for comparison). **d** RPFace-CLIP on factors seen and unseen during training, across assay, cell-line and chromosome splits, with the number of RBPs below each group. Dashed lines denote chance.

#### Normalized mass enrichment (M3)

We therefore score CLIP predictions with M3 (Eq. (6), Section 2.10), which measures where a prediction places its score mass rather than how it ranks individual nucleotides. Because any concentration of mass inside the peak counts, the footprint-only oracle that scored 0.62 AUROC and 0.50 P@*K* obtains an M3 of 1.00 (Fig. 5b). We use M3 as the primary CLIP metric and report AUROC and P@*K* alongside it.

#### Transfer from structural training

The base RPFace, Boltz-2 with and without MSAs, and AlphaFold3 all have M3 distributions centred close to chance (Fig. 5c). Fine-tuning RPFace on CLIP shifts the distribution upward, producing a positive enrichment signal that the structurally trained model lacks. Structural pretraining alone therefore does not yield detectable transfer to CLIP binding, whereas task-specific CLIP supervision enables prediction of the *in vivo* signal.

#### Generalization to unseen conditions

The CLIP-supervised signal generalizes beyond the training conditions (Fig. 5d). M3 remains positive on held-out cell lines and chromosomes, and is comparable to or higher than in the corresponding seen groups. Transfer across assays is weaker, with the unseen-assay distribution closer to chance. AUROC and P@*K* give broadly similar seen–unseen comparisons but vary much more across RBPs.

### 3.4 Ablations, partner specificity and efficiency

#### Ablation studies. To identify which components drive these results, we performed ablation studies, retraining RPFace from scratch with one component removed at a time

(Supplementary Section S9, Fig. S4). Removing the ESM2 and RNA-FM embeddings reduced AUROC for both molecules on both benchmarks. On the PDB benchmark, protein AUROC fell from 0.887 to 0.802 and RNA AUROC from 0.807 to 0.676. On the docking benchmark, they fell from 0.782 to 0.739 and from 0.644 to 0.593 (all 95% bootstrap intervals exclude zero). On the PDB benchmark the loss was largest in non-ribosomal complexes, the regime in which co-folding models are weakest (Section 3.1). The ablation of the auxiliary losses is reported in Supplementary Section S9.

#### Partner specificity

Because RPFace receives both sequences, its prediction should depend on the specific pair rather than on either molecule alone. We used groups in which one molecule binds several partners at different sites. Each prediction was scored against the true partner’s annotation and against the annotations of other partners of the same molecule. Predictions agree more strongly with the true-partner annotation on both protein and RNA, and partner swaps move the scores toward chance. The effect is strongest when the RNA is fixed and the protein partner is changed (Supplementary Section S6, Fig. S2).

#### Efficiency

RPFace trains only 2.1 million parameters, about 250 times fewer than Boltz-2 (521 million), because it builds on frozen pretrained models. We timed pairs of 308–3,016 tokens on one GPU, from sequences to per-residue predictions. RPFace was 1.6–1.9× faster than Boltz-2 with alignments and used the least memory at every size. Boltz-2 with alignments exceeded 96 GB at 3,016 tokens. On the docking benchmark, RPFace needed a median of 50 s per case from sequence alone. The docking servers have reported median turnaround times of 4–206 min, but these include server queueing and are not directly comparable [22]. The Boltz-2 embedding accounts for 69–97% of RPFace’s runtime. Removing it at inference gives a variant that runs in 10–14 s per pair with at most 6 GB of memory. On the PDB benchmark this costs little (protein AUROC 0.887 to 0.878, RNA 0.807 to 0.775). On the docking benchmark, whose complexes differ more from the training data, the cost is larger (0.782 to 0.723 and 0.644 to 0.559; Supplementary Section S7, Fig. S3).

## 4 Discussion

Protein–RNA methods are usually evaluated by a single average over data dominated by a few complex families. Our stratified benchmark shows that the interface accuracy of co-folding models on RNA is concentrated in ribosomal complexes and in the cryo-EM, lower-resolution regime that accompanies them. This accuracy collapses without MSAs or when a partner has no homologs, and docking holds only when the supplied conformations are near the bound state. RPFace predicts the binding preference of every residue and nucleotide from sequence alone and remains stable across these conditions. Its clearest advantages are on RNA interfaces, non-ribosomal complexes and partners without evolutionary information. It needs no alignment, template or structure, and runs faster and with less memory than co-folding models, which makes it suited to large-scale screening and to engineered molecules.

Beyond the model, M3 provides a peak-width-robust way to score any binding-site predictor against CLIP data, where standard ranking metrics penalize correct localization.

Several limitations remain. Cryo-EM, resolution and ribosomal identity are confounded in the available structures, so the stratified results describe one distribution shift seen from several angles. The evidence for synthetic RNAs rests on a single post-cutoff complex. When informative MSAs exist, co-folding models can still be more accurate on protein interfaces, as for DasR. Finally, *in vivo* binding remains an open challenge. Structural training alone does not transfer to CLIP, and transfer across assays stays weak after fine-tuning. The gap between solved interfaces and cellular binding is likely shaped by competition, chaperones, modifications and local concentration. Natural next steps are benchmarks that separate these factors, and models that combine sequence-level robustness with evolutionary information when it is available.

## Notes

### Competing Interest Statement

The authors have declared no competing interest.

