## Supplementary Material for "RPFace: From biased structural benchmarks to generalizable RNA–protein interface prediction"

### S1 Structural dataset construction

**PDB extraction.** We selected PDB complexes [1] containing both protein and RNA, with a reported resolution of ≤ 3.5 Å, and used biological assembly 1. For each protein–RNA chain pair, we calculated all heavy-atom distances within the complete assembly. We retained pairs with at least 25 inter-chain contacts at a distance threshold of 4.5 Å. Deduplication based on sequence and interface signatures yielded 67,716 candidate chain pairs.

**Docking benchmark.** Comparing docking methods requires complexes for which both bound and unbound structures are available. Such complexes are rare, so we used the consolidated protein–RNA docking dataset of Chong et al. [2]. It contains 235 test cases curated from PDB structures deposited up to July 2024. Each case includes separate protein and RNA input structures and a reference protein–RNA complex.

**Leakage filtering.** Filtering was performed before clustering, in four steps (Table S1). First, we removed pairs from PDB entries in the docking benchmark. Second, we used MMseqs2 [3] to exclude proteins sharing ≥ 30% sequence identity over ≥ 50% coverage with a docking-benchmark protein. Third, we searched candidate RNAs against Rfam covariance models [4] with Infernal [5], and removed RNAs assigned to a family represented in the docking benchmark. Finally, we compared protein structures with Foldseek [6] and excluded pairs whose protein matched a docking-benchmark protein with $\max(\mathrm{qTM}, \mathrm{tTM}) \geq 0.5$.

**Table S1: Structural-dataset filtering.** Counts refer to protein–RNA chain pairs.

| Stage | Removed | Remaining |
|---|---|---|
| Initial deduplicated pool | – | 67,716 |
| Protected PDB IDs | 216 | 67,500 |
| Protein sequence homology | 2,783 | 64,717 |
| Protected Rfam families | 1,716 | 63,001 |
| Protein structural homology | 9,376 | 53,625 |

**Clustering and temporal split.** After filtering, we clustered the complete remaining dataset once, obtaining 1,098 protein clusters, 635 RNA clusters and 9,029 complex clusters. A complex cluster is defined by the combination of its protein and RNA clusters. Pairs containing the same protein but different RNAs therefore belong to different complex clusters, as do pairs containing the same RNA but different proteins. This product-based definition reflects the contributions of both binding partners. The training and held-out partitions were derived from these complex clusters.

We used 1 June 2024 as the temporal cutoff, based on the initial PDB release date, because this date corresponds to the training-data cutoff of Boltz-2. Complex clusters containing both pre- and post-cutoff structures were assigned entirely to the training set, so the training set was not strictly limited to pre-cutoff structures. By contrast, clusters eligible for the held-out set contained only post-cutoff structures. No complex cluster was shared between the training and held-out sets.

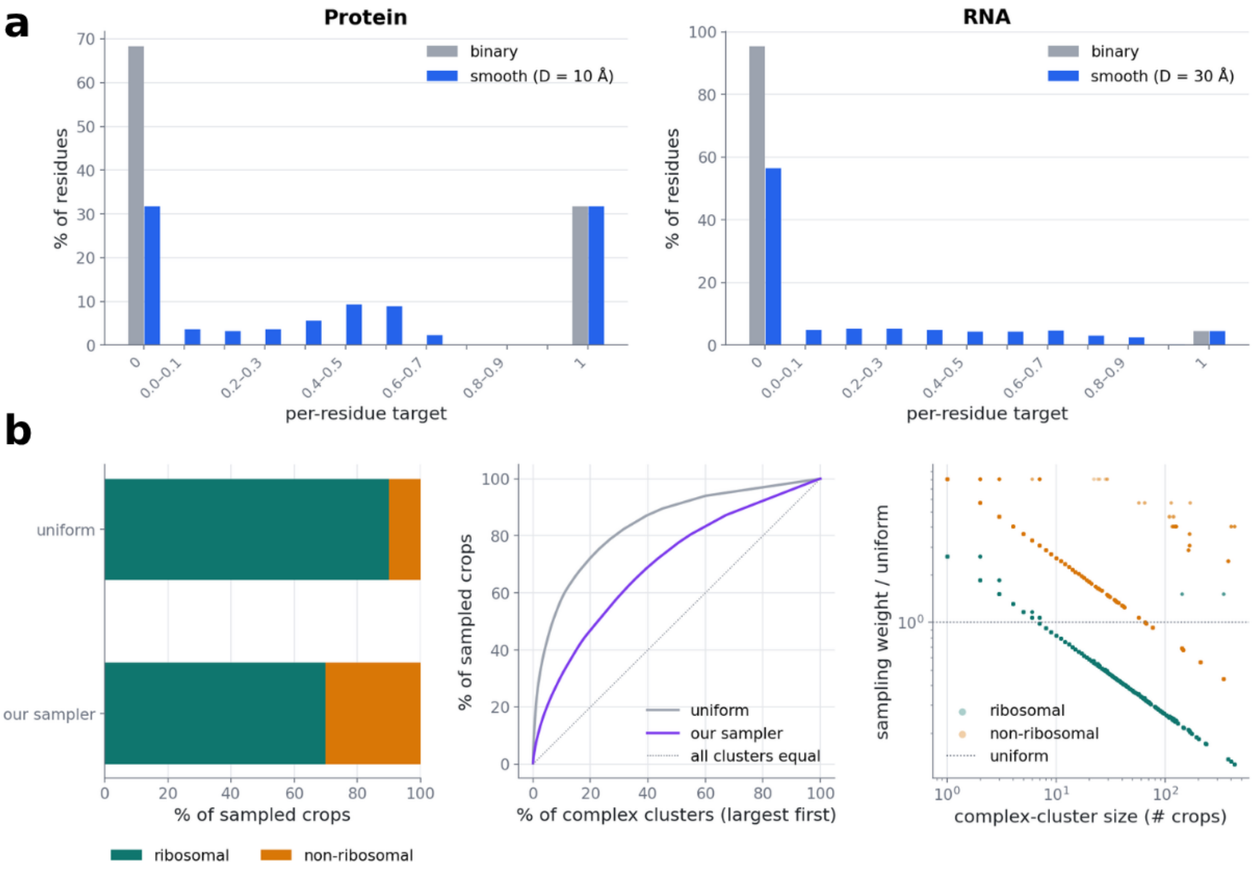


**Figure S1: Training targets and cluster-balanced sampling. a** Distributions of binary interface labels (grey) and smoothed distance-based targets (blue) for protein residues (left) and RNA nucleotides (right). Smooth targets are $t = \max(0, 1 - \delta/D)$, where $\delta$ is the distance to the nearest interface position on the same molecule, with $D = 10$ Å for proteins and $D = 30$ Å for RNA. Smoothing assigns non-zero graded targets to 36.6% of protein residues and 39.0% of RNA nucleotides. **b** Ribosomal and non-ribosomal composition before and after balanced sampling. The sampler shifts the natural 90:10 ratio to 70:30, matching the validation set. Balancing also increases the effective number of clusters from 2,575 to 5,401 and reduces the contribution of the largest 1% of clusters from 20% to 7%. Within each group, weights scale as $(1/\text{cluster size})^{0.5}$. Points outside the two dominant trends correspond to clusters containing both ribosomal and non-ribosomal crops.

**Validation set.** We constructed the validation set by randomly sampling 10–15% of the eligible complex clusters. Sampling was stratified by four factors: experimental method (X-ray crystallography or cryo-EM), ribosomal status, resolution ($< 2.5$, 2.5–3.0, or $\geq 3.0$ Å), and RNA length ($< 20$, 20–50, 50–100, or $> 100$ nucleotides). This stratification prevents the validation set from being dominated by a single class, such as ribosomal X-ray structures, and preserves the overall composition of the dataset. Validation metrics therefore reflect performance across the main structural regimes in the training distribution.

**Cluster-balanced sampling.** Training uses interface-centred crops, including spatial negatives without protein–RNA contacts. Within the ribosomal and non-ribosomal groups, crops are sampled with cluster weights proportional to $(1/\text{cluster size})^{0.5}$. The two groups are reweighted so that the natural 90:10 ratio of ribosomal to non-ribosomal crops becomes 70:30, matching the validation set. Balancing increases the effective number of clusters from 2,575 to 5,401 and reduces the contribution of the largest 1% of clusters from 20% to 7% (Fig. S1b).

### S2 CLIP dataset construction

**Sources and experimental groups.** We used the human POSTAR3 collection [7], which contains 38,361,545 peak records mapped to GRCh38. Genomic sequences were obtained from the GRCh38 primary assembly, gene annotations from GENCODE v29, and full-length RBP sequences from UniProt [8]. POSTAR3 integrates several peak-calling methods whose numerical scores represent different quantities, including $p$-values, tag counts, conversion-based scores and fold enrichment. These scores are not directly comparable, so we did not use them. Records were grouped by CLIP technology and cell line or tissue. We excluded genomic scaffolds, mitochondrial and Epstein–Barr virus contigs, non-protein-specific occupancy measurements, and proteins with fewer than 200 peaks within a group. To provide sufficient cross-protein negative evidence, each group also had to contain at least two proteins and 5,000 peaks. Sixteen of the 52 experimental groups met these criteria, spanning eCLIP, PAR-CLIP, iCLIP and HITS-CLIP.

**Coverage blocks and windows.** Within each experimental group, overlapping peaks were first merged separately for each protein. The merged peaks from all proteins were then combined into contiguous coverage blocks. Windows were sampled only from within these blocks, so every included nucleotide is covered by at least one experiment in the same experimental context. We sampled up to three windows from each coverage block, with lengths from 100 to 598 nucleotides. Each window was assigned an anchor protein. The anchor peak had to retain at least 50% of its original length after windowing and cover 10–45% of the window. When several proteins met these criteria, we selected the protein with the fewest previously assigned anchor windows. This balancing prevents highly represented proteins and densely bound genomic regions from dominating the dataset. For each window–protein pair, the nucleotide labels are

$$y_i = \begin{cases} 1, & \text{if nucleotide } i \text{ lies in a merged peak of the protein,} \\ 0, & \text{otherwise.} \end{cases} \tag{E1}$$

If a non-anchor peak was truncated to fewer than 15 nucleotides by a window boundary, we excluded the corresponding pair.

**Cross-dataset leakage control.** We jointly protected the structural and CLIP datasets, because CLIP fine-tuning was initialized from a model trained on the structural data. Protein homology searches used MMseqs2 with the same thresholds as the structural pipeline (30% identity, 50% coverage). Of the 216 CLIP RBPs, 190 had no relevant structural homology restrictions. Fourteen were homologous to proteins in the protected structural evaluation set and were excluded from CLIP training, and five were homologous to structural training proteins and were excluded from the CLIP protein holdout. The remaining seven matched proteins in both structural partitions and were removed from the CLIP dataset. The 216 proteins were independently grouped into 174 homology clusters, and each restriction was applied to every protein in the corresponding cluster. On the RNA side, we first used GENCODE annotations to remove windows overlapping structured non-coding RNA classes commonly represented in the PDB. We then applied a family-level filter using Rfam and Infernal. The annotation-based step retained 148,103 of the original 149,446 windows.

**Evaluation axes.** We defined three independent evaluation axes (Table S2). Split assignment was performed for each window–protein pair. Because the axes are not mutually exclusive, we stored a separate holdout indicator for each axis and evaluated them independently. Holding out an assay or cell line does not necessarily produce unseen RNA sequences, because the same genomic locus may occur in another retained experiment. We therefore report two versions of the assay and cell-line slices. The *strict* version is restricted to the held-out chromosomes, and the *relaxed* version has no chromosome restriction. The chromosome holdout was evaluated separately to measure generalization to unseen RNA sequences.

**Table S2: CLIP evaluation axes.** Each axis is held out independently of the others.

| Axis | Reserved data | Main question |
|---|---|---|
| Assay | PAR-CLIP, HITS-CLIP | Protocol generalization |
| Cell line | HepG2, HeLa | Cellular-context generalization |
| Chromosome | chr1, chr8 | RNA-sequence generalization |

**Fine-tuning on CLIP.** RPFace-CLIP is initialized from the structurally trained model. CLIP peaks mark crosslinked regions rather than sparse physical contacts, and the crosslink labels supervise the same RNA binding head used for structural interfaces, so the fine-tuned model adds no parameters and remains architecturally identical to the structural one. On the CLIP branch the Boltz-2 view is dropped from the fusion of Eq. (2) of the main text rather than replaced by zeros, and the pair representation is built from the sequence tracks alone, so every CLIP input is derived from sequence. RNA-FM embeddings and base-pairing probabilities for each unique window, and ESM2 embeddings for each RBP, are precomputed once with the same code that computes them during training; spot checks confirm that the cached values match those computed on the fly.

Fine-tuning keeps the structural supervision throughout. Each step draws one structural batch and one CLIP batch, forwards them separately, and takes a single optimizer step on

$$\mathcal{L}_{\mathrm{FT}} = \mathcal{L} + \lambda_{\mathrm{CLIP}}\,\mathcal{L}_{\mathrm{CLIP}}, \tag{E2}$$

where $\mathcal{L}$ is the structural objective of Eq. (E10) with the same weights. Because a single head must satisfy both label semantics, the structural term in the same step is what holds that head to the interface task. The crosslink term $\mathcal{L}_{\mathrm{CLIP}}$ is a per-nucleotide generalized cross-entropy [13] with $q = 0.7$, which scales the cross-entropy of each position by $p_y^q$ and so down-weights labels the model confidently disagrees with; this is appropriate because CLIP positives carry one-sided peak-caller noise that otherwise dominates the batch gradient once the clean data are fit. We set $\lambda_{\mathrm{CLIP}} = 0.114$, the weight at which the measured CLIP and structural gradient norms reach parity in the shared head; under an unweighted objective the CLIP term supplies about 91% of the update, and at this weight about 50%. CLIP labels exist only for the RNA, so the crosslink term is computed on nucleotides alone and the protein head receives gradient only from the structural term.

We used AdamW with a learning rate of $10^{-4}$, ten times lower than in structural training, no weight decay, and global gradient-norm clipping at 1.0, with linear warmup over the first 10% of steps followed by cosine decay. Updates with a non-finite loss or gradient norm were skipped, and an exponential moving average of the weights with decay 0.999 was maintained. CLIP batches contain 16 windows; the batch size is reduced when the product of batch size, protein length and window length exceeds $6 \times 10^6$, which bounds peak GPU memory. Retention of the structural task was monitored throughout fine-tuning as the cross-entropy and per-complex Pearson correlation of the binding heads on held-out structural crops.

### S3 Graded targets and model details

#### S3.1 Graded interface targets

We define the structural contact matrix $\boldsymbol{C} \in \{0,1\}^{P\times R}$, where $C_{ij} = 1$ if the minimum heavy-atom distance between protein residue $i$ and RNA nucleotide $j$ is at most 4.5 Å. A protein or RNA position belongs to the interface if it forms at least one such contact. Binary interface labels produce a strong class imbalance, because non-interface positions greatly outnumber interface positions. We therefore use continuous labels that decrease with spatial distance from the observed interface.

Let $\boldsymbol{x}_i^{\mathrm{prot}} \in \mathbb{R}^3$ be the mean heavy-atom coordinate of protein residue $i$ and $\boldsymbol{x}_j^{\mathrm{rna}} \in \mathbb{R}^3$ that of RNA nucleotide $j$. For each molecule separately, the distance to the nearest interface position on the same molecule is

$$\delta_i^{\mathrm{prot}} = \min_{k\in\mathcal{I}^{\mathrm{prot}}} \left\|\boldsymbol{x}_i^{\mathrm{prot}} - \boldsymbol{x}_k^{\mathrm{prot}}\right\|_2, \qquad \delta_j^{\mathrm{rna}} = \min_{l\in\mathcal{I}^{\mathrm{rna}}} \left\|\boldsymbol{x}_j^{\mathrm{rna}} - \boldsymbol{x}_l^{\mathrm{rna}}\right\|_2, \tag{E3}$$

where $\mathcal{I}^{\mathrm{prot}}$ and $\mathcal{I}^{\mathrm{rna}}$ are the protein and RNA interface positions, so $\delta = 0$ on the interface. The labels are

$$t_i^{\mathrm{prot}} = \max\left(0,\ 1 - \delta_i^{\mathrm{prot}}/D_{\mathrm{prot}}\right), \qquad t_j^{\mathrm{rna}} = \max\left(0,\ 1 - \delta_j^{\mathrm{rna}}/D_{\mathrm{rna}}\right), \tag{E4}$$

with $D_{\mathrm{prot}} = 10$ Å and $D_{\mathrm{rna}} = 30$ Å. Spatial negative complexes, which contain no protein–RNA contacts, receive $\boldsymbol{t} \equiv 0$ for both molecules. The resulting target distributions are shown in Fig. S1a.

**Why the scale is absolute.** An earlier version used the same construction but normalized distances per structure, $\tilde{t}_i^{\mathrm{prot}} = 1 - \delta_i^{\mathrm{prot}}/\max_k \delta_k^{\mathrm{prot}}$. This may suit complete structures, but not the interface-centred crops used here. In a crop, $\max_k \delta_k^{\mathrm{prot}}$ reflects the spatial extent of the crop rather than a physical property of the binding site. Because each crop is centred on an interface, the farthest residue always receives 0, whether it is 8 Å or 40 Å from the interface. The labels then depend on crop geometry and are not comparable across complexes. This normalization also shifts the labels upward: the mean protein-side label was 0.72. The fixed cutoffs of Eq. (E4) remove this dependence. Each target represents an absolute distance from the interface on a consistent physical scale.

**The output is a censored distance.** Equation (E4) is an affine function of $\delta$ truncated to the unit interval. Predictions in the uncensored range can therefore be converted directly to distances:

$$\hat{\delta}_i^{\text{prot}} = D_{\text{prot}}\big(1 - \hat{m}_i^{\text{prot}}\big), \qquad \hat{\delta}_j^{\text{rna}} = D_{\text{rna}}\big(1 - \hat{m}_j^{\text{rna}}\big). \tag{E5}$$

The model thus performs censored regression on the distance to the interface through a logistic link. The endpoints carry limited distance information: a score of 1 indicates an interface position, and a score of 0 a distance of at least $D$. Intermediate scores correspond to distances within the physical range. This is a stronger and more directly testable prediction than a ranking of binding preferences.

**Class imbalance.** The protein interface fraction is $\pi^{\text{prot}} = P^{-1} \sum_i y_i^{\text{prot}}$, and $\pi^{\text{prot}}$ and its RNA counterpart vary widely across complexes. Both have heavy-tailed, strongly bimodal distributions. For example, a ribosomal protein embedded in rRNA can have $\pi^{\text{prot}} \approx 0.4$, whereas a protein bound to a long RNA transcript can have $\pi^{\text{rna}} < 0.03$. A single class-reweighting constant is therefore unsuitable for all complexes. Pooling complexes with different interface fractions also combines prediction tasks with very different class balances. This heterogeneity is accompanied by substantial imbalance in the composition of the training set (Fig. S1b), which has two consequences. First, the training objective combines position-level terms, which are dominated by longer chains, with per-complex terms that weight complexes independently of chain length. Second, each precision-based metric is interpreted relative to the empirical interface fraction of its stratum.

#### S3.2 Input representations and gated fusion

Each protein residue or RNA nucleotide is represented by three complementary views, combined through a learned convex gate. For proteins these are the Boltz-2 trunk representation $\boldsymbol{s}^{\text{prot}} \in \mathbb{R}^{P\times d_s}$ with $d_s = 384$ [9], the frozen ESM2 representation $\boldsymbol{e} \in \mathbb{R}^{P\times 1280}$ [10], and a physicochemical prior $\boldsymbol{q}^{\text{prot}}$. For RNA they are the Boltz-2 representation $\boldsymbol{s}^{\text{rna}} \in \mathbb{R}^{R\times d_s}$, the frozen RNA-FM representation $\boldsymbol{g} \in \mathbb{R}^{R\times 640}$ [11], and a nucleotide prior $\boldsymbol{q}^{\text{rna}}$. Boltz-2 representations are precomputed in single-sequence mode with an empty MSA. Neither the Boltz-2 pair representation nor predicted coordinates are used. For a protein or RNA token $a_i$, the prior is

$$q_i = \psi\big([\,\text{Emb}(a_i);\ \boldsymbol{\phi}(a_i);\ \text{PE}(i)\,]\big), \tag{E6}$$

where Emb is a learned 32-dimensional token-identity embedding, $\boldsymbol{\phi}$ collects physicochemical descriptors (hydropathy, formal charge, aromaticity and residue volume for proteins; a purine indicator and hydrogen-bond donor and acceptor counts for RNA), PE is a 16-dimensional sinusoidal positional encoding, and $\psi$ is a two-layer perceptron. All descriptors are standardized with fixed population constants rather than batch statistics, so the encoding of a token is independent of the complex in which it occurs.

The three views are projected to a common dimension $d = 96$ by layer-normalized affine maps, giving $h_{1,i}$ (Boltz-2), $h_{2,i}$ (language model) and $h_{0,i}$ (prior). These are fused as in Eq. (2) of the main text. The prior enters as an ungated residual, so the gate cannot remove it. During training, the projected Boltz-2 view is dropped with probability $p_{\text{drop}} = 0.1$, provided at least one view remains. Dropout is applied after the projection rather than by feeding a zero input to it. For CLIP fine-tuning, the Boltz-2 view is removed and the model relies on the language-model and prior views.

#### S3.3 Ensemble-averaged secondary-structure propagation

Base-pairing probabilities are computed with ViennaRNA [12]. The propagation of Eq. (3) in the main text integrates information across the Boltzmann ensemble rather than using only the single partner predicted by a minimum-free-energy structure. This is appropriate because RNA molecules occupy multiple conformations, and it does not depend on a particular crystallographic conformer. Eq. (3) is the only message-passing operation retained in the final model. Ablation experiments supported removing three other propagation schemes: protein local-sequence propagation, RNA local-sequence propagation, and propagation based on RNA-FM embedding similarity.

#### S3.4 Partner coupling and multi-scale mixing

The protein and RNA representations are coupled through bidirectional cross-attention with $H = 4$ heads. Each protein position incorporates information from the RNA partner, and each RNA position from the protein partner. A compact pair representation is updated from both sequence tracks. Let $h_{\text{loc}}$ and $h_{\text{crs}}$ be the representations before and after cross-attention. A per-token convex gate $\beta_i$ combines them, allowing the model to reduce the contribution of partner-derived information when it is uninformative.

Both tracks are then processed by a dilated convolutional stack with dilation factors $(1, 2, 4, 8, 16, 32)$, kernel width 5, gated linear unit activations and residual connections. Every layer is masked to exclude padded positions. The receptive field spans 253 positions (±126 around each token). This range captures extended contiguous binding regions without imposing a fixed input window. Multi-scale mixing at this stage supports transfer from interface-centred training crops to full-length chains at inference, and reduces sensitivity to sequence length.

### S4 Training objective

**Principal term.** Because the targets are continuous, the binding loss (Eq. (4) of the main text) is a cross-entropy against a target distribution, not a Bernoulli log-likelihood for an observed binary outcome. It decomposes as

$$\ell(z, t) = \mathbb{H}(t) + \mathrm{KL}\big(\mathrm{Ber}(t) \,\|\, \mathrm{Ber}(\sigma(z))\big). \tag{E7}$$

The unique minimum is at $\sigma(z) = t$, so the loss is proper for recovering the target profile, but its attainable minimum is the target entropy $\mathbb{H}(t)$ rather than zero. The targets form a mixed distribution on $[0, 1]$: interface positions give a point mass at 1, positions beyond the cutoff $D$ give a point mass at 0, and positions in the surrounding rim take continuous values between these endpoints. Because $\mathbb{H}(0) = \mathbb{H}(1) = 0$, only rim positions contribute to the irreducible loss, in proportion to the rim fraction of each complex. Since $D_{\mathrm{rna}} = 3D_{\mathrm{prot}}$, RNA targets generally contain a larger graded region and have a higher loss floor. The protein and RNA terms are therefore not directly comparable, and their relative magnitudes should not be read as differences in task difficulty. Each term is normalized over valid positions pooled across the batch. This micro-average deliberately gives more weight to long RNA chains, where nucleotide-level prediction is hardest. The overlap term below is normalized per sequence and acts as a counterweight.

**Overlap term.** Cross-entropy treats positions independently and does not constrain the shape or overlap of the predicted interface. We therefore add a Lovász hinge loss, a tight convex extension of the Jaccard loss that directly optimizes set-level agreement. It is computed per sequence:

$$\mathcal{L}_{\mathrm{lov}} = \frac{1}{B}\sum_{n=1}^{B}\sum_{k}\mathrm{ReLU}\big(1 - z_{\pi(k)} s_{\pi(k)}\big)\, g_k, \qquad s = 2\bar{y} - 1, \tag{E8}$$

where $\bar{y}$ are the binary interface labels, $\pi$ orders the margin errors from largest to smallest, and $g_k$ is the discrete change in the Jaccard index along this ordering. Unlike cross-entropy, this term is normalized per sequence, which reduces the dominance of long chains. Soft-Dice and focal-Tversky losses, which also support graded targets, were implemented as alternatives but not used in the reported model.

**Auxiliary supervision.** We supervise the pair representation with $\mathcal{L}_{\mathrm{pair}} = \mathrm{BCE}(\mathrm{PairHead}(\mathbf{Z}), \boldsymbol{\Gamma})$. Here $\boldsymbol{\Gamma}$ is a graded contact map constructed as in Eq. (E4), but using distances in the joint centroid space of the complex. We also supervise a symmetrized bilinear prediction from the RNA track against the true base-pairing matrix ($\mathcal{L}_{\mathrm{ss}}$), and use a three-state classification loss for protein secondary structure ($\mathcal{L}_{\mathrm{sse}}$).

Two auxiliary losses reduce shortcut learning during view fusion. Single-view prediction heads are attached to each projected protein and RNA view ($\mathcal{L}_{\mathrm{view}}$). Additional heads are attached to the representations before interaction with the partner ($\mathcal{L}_{\mathrm{pre}}$). All these heads are trained against the same graded binding targets. Together with view dropout, they are intended to prevent the fusion gate from collapsing onto the view that is easiest to optimize early in training; their measured effect on the gate and on accuracy is reported in Section S9. We further regularize the average use of the learned views with

$$\mathcal{L}_{\mathrm{bal}} = \big\|\mathbb{E}_i[\alpha_i] - \tfrac{1}{2}\mathbf{1}\big\|_2^2. \tag{E9}$$

This constrains only the batch-averaged gate weights. Individual tokens may still assign all weight to one view, while the marginal use of the two views stays balanced.

**Total objective.**

$$\begin{aligned}\mathcal{L} = {} & \mathcal{L}_{\mathrm{bind}} + \lambda_{\mathrm{lov}}\mathcal{L}_{\mathrm{lov}} + \lambda_{\mathrm{pair}}\mathcal{L}_{\mathrm{pair}} + \lambda_{\mathrm{ss}}\mathcal{L}_{\mathrm{ss}} \\ & + \lambda_{\mathrm{sse}}\mathcal{L}_{\mathrm{sse}} + \lambda_{\mathrm{view}}\mathcal{L}_{\mathrm{view}} + \lambda_{\mathrm{pre}}\mathcal{L}_{\mathrm{pre}} + \lambda_{\mathrm{bal}}\mathcal{L}_{\mathrm{bal}},\end{aligned} \tag{E10}$$

with $\lambda_{\mathrm{pair}} = 0.1$, $\lambda_{\mathrm{ss}} = 1.0$, $\lambda_{\mathrm{sse}} = 0.2$, $\lambda_{\mathrm{view}} = 0.2$, $\lambda_{\mathrm{pre}} = 0.1$, $\lambda_{\mathrm{bal}} = 0.01$ and $\lambda_{\mathrm{lov}} = 0.1$. The Lovász weight was selected on the validation set; all other weights were fixed before training. The structural model was trained with

AdamW (learning rate $10^{-3}$, weight decay $10^{-2}$), global gradient-norm clipping at 1.0, linear warmup over the first 10% of steps followed by cosine decay, view dropout with probability 0.1, and an exponential moving average of the weights with decay 0.999, for 200 epochs with random seed 0.

### S5 Evaluation metrics

**Ranking metrics.** Because RPFace outputs continuous preferences, we evaluate ranking quality rather than thresholded contacts. Experimental contacts define only the reference positives. For comparisons independent of RPFace's scoring function, structure-based methods are scored from predicted coordinates as $s = 1/d$, where $d$ is the minimum distance from each residue or nucleotide to the other molecule. All methods thus produce continuous rankings without method-specific contact thresholds. AUROC measures ranking across positive and negative positions and does not depend on class prevalence. AUPRC, Precision@$K$ and Success@5 emphasize top-ranked predictions and depend on the positive fraction, so each is interpreted relative to the interface fraction of its stratum. Precision@$K$ is the fraction of true positives among the $K$ top-ranked positions, with $K$ equal to the number of positive positions in the molecule, so that a perfect ranking scores 1 and Recall@$K$ coincides with Precision@$K$. Success@5 is 1 if at least one of the five top-ranked positions is a true positive and 0 otherwise, averaged over molecules.

**AUROC ceiling under broad peaks.** A CLIP peak represents a crosslinked RNA fragment and is generally wider than the physical protein footprint. Suppose a called peak has width $m$ and the true footprint has width $w \le m$. Consider an oracle that assigns a high score exactly to the footprint and a uniform low score elsewhere. With the whole peak as the positive label, footprint positions outrank every negative, and the remaining peak positions tie with the negatives. Hence

$$\mathrm{AUROC}_{\mathrm{oracle}} = \frac{w}{m} + \frac{1}{2}\left(1 - \frac{w}{m}\right) = \frac{1}{2}\left(1 + \frac{w}{m}\right). \tag{E11}$$

A biologically precise predictor can therefore obtain a modest nucleotide AUROC simply because the annotation is wider than the footprint (main text Fig. 5a,b). Precision@$K$, Recall@$2K$ and Success@5 are retained for the CLIP data because they answer a different question, namely whether the top of the ranking lies on the peak, but they are limited by the same dilation: a correct $w$-nucleotide footprint can occupy only $w$ of the $m$ labelled positions, and the remainder of the top $K$ is filled by tie-breaking. We therefore measure the attainable ceiling for each of these metrics with the footprint oracle rather than assuming it to be 1.

**Normalized mass enrichment (M3).** Let $m = \sum_i y_i$ and $p = m/L$. We remove the bottom of each score profile using its fifth percentile,

$$\tilde{s}_i = \max\bigl(s_i - q_{05}(s), 0\bigr), \tag{E12}$$

and measure the fraction of the remaining score mass in labelled positions:

$$E = \frac{1}{p}\,\frac{\sum_i y_i \tilde{s}_i}{\sum_i \tilde{s}_i}. \tag{E13}$$

If score mass is independent of the labels, the expected mass fraction inside a peak is $p$, and hence $E = 1$. If all score mass is inside the peak, $E = 1/p$. We therefore normalize as

$$E_{\mathrm{norm}} = \frac{E - 1}{1/p - 1}, \tag{E14}$$

which maps the chance expectation to 0 and perfect concentration inside the peak to 1. A value converts to an absolute enrichment through $p$: the share of score mass inside the peak is $p + E_{\mathrm{norm}}(1 - p)$. At the median coverage of this split ($p = 0.29$), a value of 0.1 corresponds to 36% of the mass inside the peak, compared with 29% at chance.

**Properties of M3.** Let $s'_i = a s_i + b$ with $a > 0$. Then $q_{05}(s') = a\, q_{05}(s) + b$, so $\tilde{s}'_i = a\tilde{s}_i$ and $E$ is unchanged; M3 is therefore invariant to positive affine transformations of the scores. Because $0 \le \sum_i y_i \tilde{s}_i \le \sum_i \tilde{s}_i$, we have $E \in [0, 1/p]$ and M3 $\in [-p/(1-p),\ 1]$. M3 equals 1 exactly when no baseline-corrected mass lies outside the peak. This holds for the footprint oracle whatever the footprint width, provided the footprint lies inside the peak and covers less than 95% of the window, so that $q_{05}$ equals the low score. If score mass is independent of the labels, the expected share inside the peak is $p$, so $E = 1$ and M3 $= 0$ in expectation. M3 is undefined when $p \in \{0, 1\}$ or when the profile is constant; such cases are excluded.

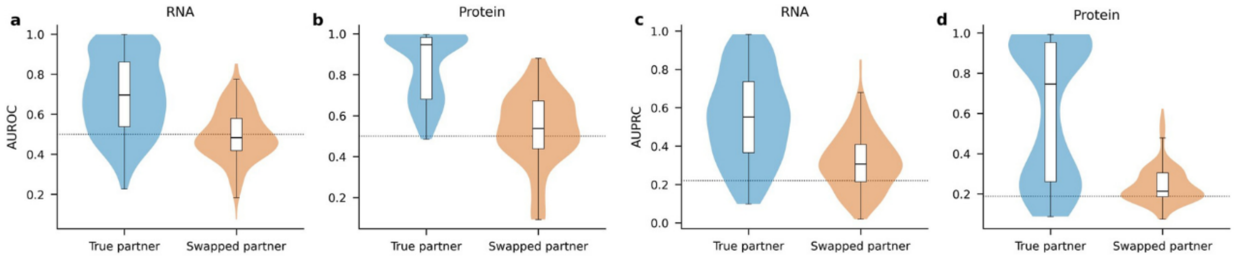


**Figure S2: RPFace predictions are specific to the binding partner.** **a** RNA AUROC. **b** Protein AUROC. **c** RNA AUPRC. **d** Protein AUPRC. Groups contain one molecule interacting with at least two partners at different sites. True-partner scores use the matching annotation; swapped-partner scores use an annotation from another partner of the same molecule. Violins show group-level distributions, and box plots show medians and interquartile ranges. Dotted lines indicate random expectation.

**Reference arms and uncertainty on the CLIP data.** Chance is measured rather than assumed. For M3 we use a circular-shift null, which preserves each score profile's marginal distribution and autocorrelation exactly and destroys only its alignment to the label; for the ranking metrics we use independent noise. We additionally report a positional baseline that scores only proximity to the centre of the window, because window construction leaves a residual centring bias that a model must clear to have demonstrated sequence-specific signal. All intervals are cluster bootstraps over RBP rather than over windows, since windows from one protein are correlated, and comparisons between methods use the paired per-RBP difference rather than overlapping marginal intervals.

### S6 Partner specificity

RPFace receives both the protein and the RNA sequence, so its prediction should depend on the specific pair rather than on either molecule alone. We tested this using groups in which one molecule interacts with several partners at different binding sites. For each prediction, we compared the score against the annotation of the true partner and against annotations from other partners of the same molecule. A model that ignores the partner would perform similarly in both cases. Predictions agree more strongly with the true-partner annotation on both protein and RNA, while partner swaps move the scores toward chance (Fig. S2). The effect is stronger when the RNA is fixed and the protein partner is changed.

### S7 Runtime and the contribution of the Boltz-2 embedding

**Measurement protocol.** Runtime was measured on five validation pairs chosen to span complex sizes (308–3,016 tokens), on a single otherwise idle NVIDIA RTX PRO 6000 Blackwell GPU (96 GB). Each pipeline stage ran as a fresh process, so model loading is included. Each configuration was repeated three times after a discarded warm-up run, and we report medians. Peak GPU memory was sampled every 100 ms. RPFace was timed in two stages: first the Boltz-2 trunk pass in single-sequence mode with three recycles, and then a stage that loads RPFace, ESM2 and RNA-FM, computes the language-model embeddings and base-pair probabilities with empty feature caches, and runs the forward pass. The runtime of RPFace without the Boltz-2 embedding is that of the second stage, since dropping the Boltz-2 branch does not change its computation. Boltz-2 [9] was run with three recycles, 200 sampling steps and one diffusion sample. Alignments came from the ColabFold MMseqs2 server, which returns stored results for previously searched sequences. To time a genuine search, we introduced a single substitution at a different position in each repeat.

**Runtime and memory.** RPFace was 1.6–1.9× faster than Boltz-2 with alignments (Fig. S3c), for which the alignment search alone accounted for 25–54% of the total runtime. Compared with Boltz-2 without alignments, RPFace was slightly slower below about 1,000 tokens, owing to model loading, and 1.1–1.3× faster above it. RPFace also had the lowest peak memory at every size (Fig. S3b). Boltz-2 with alignments exceeded the 96 GB of GPU memory at 3,016 tokens in all three repeats.

**AlphaFold3.** We could not time AlphaFold3 [14] under the same conditions. We accessed it only through its web server, whose runtimes include queueing on undisclosed hardware. For reference, its published single-GPU inference times are

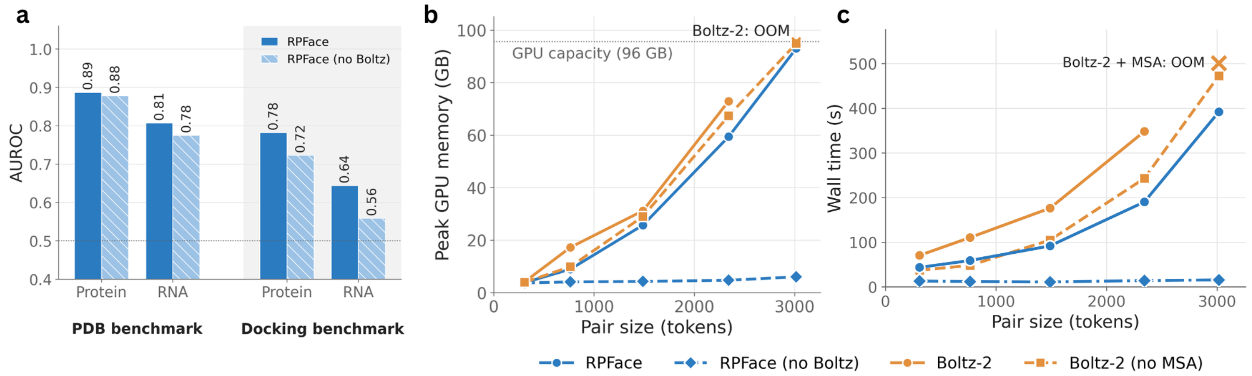


**Figure S3: Contribution of the Boltz-2 embedding and computational cost of RPFace.** **a** AUROC of RPFace (solid blue) and RPFace without the Boltz-2 embedding (light blue, hatched) on the PDB benchmark and the docking benchmark (grey background), for protein residues and RNA nucleotides. The dotted line marks random expectation (0.5). **b** Peak GPU memory over all steps of each pipeline. **c** End-to-end wall time per protein–RNA pair, from sequences to per-residue predictions, including model loading (median of three repeats). In **b** and **c**, pair size is the number of protein residues plus RNA nucleotides. Blue marks RPFace (circles) and RPFace without the Boltz-2 embedding (diamonds). Orange marks Boltz-2 with an MSA (circles) and without one (squares). The RPFace time includes computing the Boltz-2 embedding (trunk only, without MSA). The time of Boltz-2 with MSA includes an uncached ColabFold MSA search. Boltz-2 used 3 recycling steps, 200 sampling steps and one sample; with MSA, it ran out of memory (OOM) at 3,016 tokens in all three repeats. All runs used one NVIDIA RTX PRO 6000 GPU (96 GB).

34, 144 and 367 s at 1,024, 2,048 and 3,072 tokens on an NVIDIA H100 [15]. These timings use 10 trunk recycles and five diffusion samples [14]. They cover only the network itself, excluding compilation and the data pipeline. That pipeline runs jackhmmer and nhmmer searches over seven protein and RNA sequence databases plus a template search [14]. It accounts for most of the end-to-end runtime of AlphaFold3 on complex inputs, and its memory requirement grows steeply with RNA length [16]. The inference step of AlphaFold3 alone is therefore comparable in time to the complete RPFace pipeline, which requires no alignment or template search at any stage.

**Docking servers.** On the docking benchmark, Chong et al. reported median turnaround times per case of 4 min for ZDOCK, 17 min for MDockPP, 30 min for LightDock, 70 min for HDOCK and 206 min for ClusPro [2]. These times include queueing on shared web servers of unspecified hardware. They also exclude obtaining the unbound input structures that docking requires, so they are not directly comparable to our local measurements. On the same 235 cases, RPFace needed a median of 50 s per case, starting from sequences alone.

**Removing the Boltz-2 embedding.** RPFace trains only 2.1 million parameters, about 250× fewer than Boltz-2 (521 million), because it builds on frozen pretrained models. Its runtime is dominated by the Boltz-2 embedding, which takes 69–97% of the total; the rest of the pipeline takes only seconds. We therefore evaluated the same trained model with this embedding removed at inference. Without it, RPFace ran in 10–14 s per pair at every size, 39× faster than Boltz-2 without alignments at 3,016 tokens, and used at most 6 GB of GPU memory (Fig. S3b,c).

On the PDB benchmark, accuracy changed only slightly (Fig. S3a): protein AUROC went from 0.887 to 0.878, and RNA AUROC from 0.807 to 0.775. These values remain comparable to AlphaFold3 with alignments (0.880 and 0.767) and well above both co-folding models without alignments. On the docking benchmark, the loss was larger: protein AUROC fell from 0.782 to 0.723 and RNA AUROC from 0.644 to 0.559. The Boltz-2 embedding therefore contributes little on the PDB benchmark but substantially on the docking benchmark, whose complexes differ more from the training data. Without it, RPFace becomes an alignment-free predictor that runs in seconds on a single GPU, suited to large-scale screening when a modest loss of accuracy is acceptable.

### S8 Comparison with ProRB on FoldBench

We compared RPFace with ProRB [17], a recent sequence-based method, on the nine FoldBench [18] protein–RNA targets that Xue et al. used to compare ProRB with AlphaFold3. We used the same targets, chains and binding-site definition as Xue et al., and took the ProRB and AlphaFold3 values from their publication.

FoldBench limits homology to structures deposited before January 2023. Both RPFace and ProRB were trained on later data, so we quantified the similarity of each target to each method's training data. We applied the criterion used to construct the RPFace training data: a target is independent of a training set only if no training protein shares at least 30% sequence identity over at least 50% coverage (MMseqs2), no training RNA belongs to the same Rfam family, and its PDB entry is not part of the training set. Against Q-BioLiP, the source of the ProRB training data, all nine targets fail this criterion: each target protein has a counterpart with 97–100% sequence identity. Against the RPFace training set, five targets fail and four are independent, with no detectable protein similarity. Xue et al. excluded six FoldBench targets that overlapped with the ProRB training set, but did not report the similarity of the remaining nine to it.

On the four targets independent of its training data, RPFace reached a mean AUROC of 0.708 for protein residues and 0.715 for RNA nucleotides. Xue et al. report 0.575 and 0.544 for ProRB and 0.557 and 0.643 for AlphaFold3; the published values are averages over all nine targets. On all nine targets, RPFace reached 0.786 and 0.611.

### S9 Ablation studies

**Protocol.** We ablated two components of RPFace: the language-model views (ESM2 for proteins, RNA-FM for RNA) and the auxiliary losses. Each variant was retrained from scratch with the recipe of Section S4 (200 epochs, seed 0) and exactly one change. All models, including the reference, were evaluated with their raw weights at epoch 200. Without the language-model views, the fusion of Eq. (2) of the main text combines only the Boltz-2 view and the prior. Without the auxiliary losses, , $\lambda_{\text{pair}} = \lambda_{\text{ss}} = \lambda_{\text{sse}} = \lambda_{\text{view}} = \lambda_{\text{pre}} = 0$, and all other terms of Eq. (E10) are unchanged. We evaluated on the 233-pair PDB benchmark and the 235-case docking benchmark. Differences are reported as without minus with, with 95% bootstrap intervals that resample clusters on the PDB benchmark and complexes on the docking benchmark. Intervals that exclude zero are marked as significant.

**Language-model views.** Removing ESM2 and RNA-FM lowered AUROC in all four comparisons (Fig. S4a). On the PDB benchmark, AUROC fell from 0.887 to 0.802 for proteins and from 0.807 to 0.676 for RNA. On the docking benchmark, it fell from 0.782 to 0.739 and from 0.644 to 0.593. AUPRC fell in three of the four comparisons; the difference for RNA on the docking benchmark was not significant. On the PDB benchmark, the loss occurred in both ribosomal and non-ribosomal complexes and was largest in the non-ribosomal group (Fig. S4b). On the docking benchmark, it was significant for easy targets on both molecules and for medium targets on proteins. The intervals for medium RNA targets and for both molecules on hard targets include zero.

Removing the Boltz-2 embedding cost much less on the PDB benchmark: 0.009 protein and 0.032 RNA AUROC (Section S7). That experiment removed the view at inference from the trained model, whereas the language-model ablation was retrained, so the two are not strictly comparable. Both nevertheless indicate that the language-model views, not the Boltz-2 embedding, carry most of the accuracy on this benchmark.

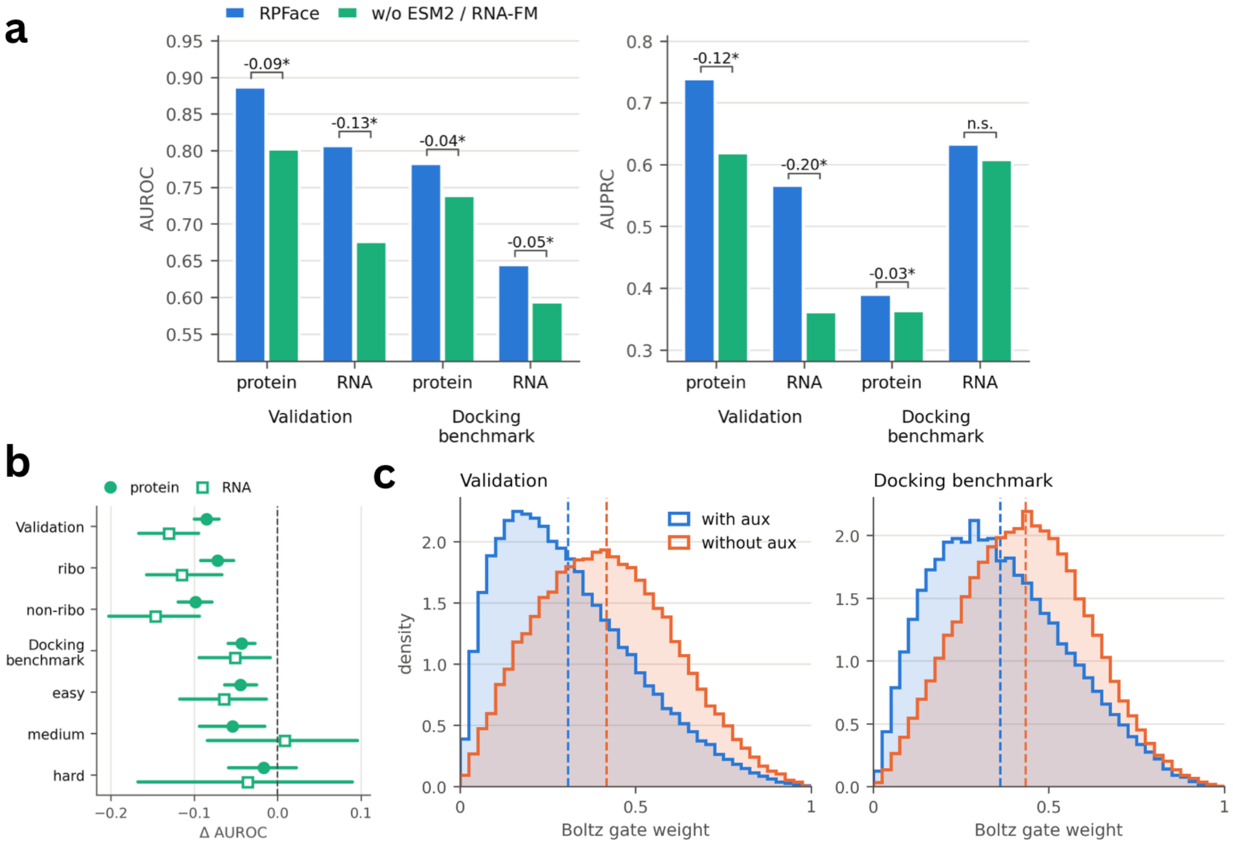


**Figure S4: Ablation of the language-model views and the auxiliary losses.** **a** Mean per-complex AUROC (left) and AUPRC (right) of RPFace and of the same model retrained without the ESM2 and RNA-FM embeddings, for protein residues and RNA nucleotides on the PDB benchmark and the docking benchmark. Brackets give the difference (without minus with). $*$, 95% bootstrap interval excludes zero; n.s., not significant. **b** AUROC difference with 95% interval on the PDB benchmark (overall, ribosomal, non-ribosomal) and on the docking benchmark (overall and by difficulty; easy, medium and hard are the Easy, Medium and Challenging classes of Chong et al. [2]). Filled circles, protein; open squares, RNA. The dashed line marks no change. **c** Distribution of the per-residue weight that the protein fusion gate assigns to the Boltz-2 view, for RPFace with and without the auxiliary losses. The ESM2 view receives the complement, so 0 means ESM2 only and 1 means Boltz-2 only. Dashed lines mark the means.
